# Coevolution of reproducers and replicators at the origin of life and the conditions for the origin of genomes

**DOI:** 10.1101/2022.09.22.509036

**Authors:** Sanasar G. Babajanyan, Yuri I. Wolf, Andranik Khachatryan, Armen Allahverdyan, Purificacion Lopez-Garcia, Eugene V. Koonin

## Abstract

There are two fundamentally distinct but inextricably linked types of biological evolutionary units, reproducers and replicators. Reproducers are cells and organelles that reproduce via various forms of division and maintain the physical continuity of compartments and their content. Replicators are genetic elements (GE), including genomes of cellular organisms and various autonomous elements, that both cooperate with reproducers and rely on the latter for replication. All known cells and organisms comprise a union between replicators and reproducers. We explore a model in which cells emerged via symbiosis between primordial ‘metabolic’ reproducers (protocells) which evolved, on short time scales, via a primitive form of selection and random drift, and mutualist replicators. Mathematical modeling identifies the conditions, under which GE-carrying protocells can outcompete GE-less ones, taking into account that, from the earliest stages of evolution, replicators split into mutualists and parasites. Analysis of the model shows that, for the GE-containing protocells to win the competition and to be fixed in evolution, it is essential that the birth-death process of the GE is coordinated with the rate of protocell division. At the early stages of evolution, random, high-variance cell division is advantageous compared to symmetrical division because the former provides for the emergence of protocells containing only mutualists, preventing takeover by parasites. These findings illuminate the likely order of key events on the evolutionary route from protocells to cells that involved the origin of genomes, symmetrical cell division and anti-parasite defense systems.

**Significance:** The origin of life, which is equivalent to the origin of cells, is arguably the greatest enigma in biology. The remarkable complexity characteristic of even the simplest extant cells could only evolve from simpler, pre-biological entities. Reconstructing that pre-cellular stage of evolution is a hard challenge. We present an evolutionary scenario in which cells evolved via symbiosis between protocells that harbored protometabolic reaction networks, could divide and were subject to selection, but lacked genomes, and primordial genetic elements. Mathematical modeling reveals conditions for the survival of such symbionts and the origin of modern-type genomes, in particular, coordination of the rates of protocell division and replication of genetic elements as well as random division of protocells.

## Introduction

Replication of genetic information is naturally considered a fundamental – or, often, the central – property of evolving biological entities, both cellular organisms and genetic parasites. All these entities possess genomes that are often also called replicators (1–3). Evidently, however, life is not limited to information transmission. An adequate supply of energy and building blocks, which depends on spatial compartmentalization, is essential for the evolution of replicators. Hence the fundamental split of all propagating biological entities into reproducers and replicators (3). Cells are reproducers: their propagation is not limited to the replication of the genome but rather, involves reproduction of the entire cellular organization that provides the niche for the replicators. Although the genome carries the instructions for the production of all cell components, it is in itself insufficient for reproduction: *Omnis cellula e cellula*. The entire evolutionary history of life is an uninterrupted, physically continuous tree of cell divisions (in which the dead branches, evidently, overwhelmingly outnumber the growing ones). By contrast, all diverse genetic elements (GE) including both cellular and organellar genomes and genetic parasites (viruses, transposons) are replicators that recruit cellular molecular machinery for some of the key functions required for their replication, in particular, translation of the GE genes (4).

The extant reproducers (cells) that necessarily host a mutualistic replicator (the genome) are compartments bounded by phospholipid membranes permeated by diverse proteins containing hydrophobic transmembrane segments (5). Compartmentalization is essential not only for preventing diffusion of small molecules into the environment and thus keeping their concentrations inside the reproducer at levels sufficient to sustain metabolic reaction networks as well as replication, but also to maintain the integrity of selectable units that consist of reproducers together with the replicators inside them. The membranes perform essential transport functions, that is, selectively import molecules and ions that are required for cell reproduction (including replication), and expel toxic molecules and ions, often in an energy-dependent manner. In respiring cells, membranes also harness the energy released in oxidation reactions to produce ion gradients that are then transformed into the energy of the macroergic phosphodiester bond of ATP. Replicators lack such active, energizable membranes and thus depend on the reproducers for energy and building blocks required for replication. Furthermore, replicators and reproducers dramatically differ in terms of the chemistries involved in their propagation. The replication process requires only narrowly focused chemistry, namely, nucleotide polymerization. Evidently, this process is underpinned by the far more complex reactions of nucleotide biosynthesis, but these are supplied by the reproducer. In contrast, even the simplest reproducers exercise a rich repertoire of chemical reactions, with at least 1000 distinct small molecules metabolized by any cell type (6).

All replicators are hosted by reproducers and depend on the hosts for energy and building blocks. However, in terms of their relationships with the host reproducer, replicators span the entire range from (near) full cooperativity and a mutualistic relationship with the host reproducer in the case of cellular genomes through commensalism in the case of plasmids and transposons, to aggressive parasitism, in the case of lytic viruses (7, 8). Arguably, only a mutualistic, obligatory union of a host reproducer with a resident replicator(s), the genome that carries instructions for the reproduction of the host, can be considered a life form (organism), in the crucial sense of temporally continuous, robust reproduction and the ensuing evolutionary autonomy (9, 10){Ruiz-Mirazo, 2004 #3034}. Thus, life took off when replicators evolved to encode components of the host reproducers, providing for the long-term persistence and evolution of the latter.

Origin of life often has been discussed in terms of competing ‘metabolism first’ vs ‘information first’ scenarios (11), which can be reformulated as “reproducers first or replicators first?” The question seems formidable and resembles a chicken and egg problem. Indeed, the fundamental differences between reproducers and replicators notwithstanding, these two types of evolving biological entities are inextricably linked. To the best of our knowledge, there are no ‘pure’ reproducers in the extant biosphere: all modern cells as well as some organelles, such as mitochondria and chloroplasts, comprise an obligatory mutualistic union of a reproducer and a replicator(s). We submit, however, that this, currently essential relationship did not exist at the primordial, pre-biological stage of evolution, which started with primitive reproducers and eventually led to the emergence of the mutualistic reproducer-replicator systems. Indeed, emergence of replication is inconceivable without a steady supply of energy and building blocks, which only can be provided by a proto-metabolism with sufficient temporal stability, that is, by some form of primordial reproducers (12, 13). Realistically, pre-biological evolution must have started with reproducers that initially did not carry any replicators within them, but rather comprised self-sustaining proto-metabolic circuits confined within membrane vesicles (14, 15). Reconstruction of these primordial reaction networks is a separate, challenging task that has been attempted in many studies (16–18) and is beyond the scope of this work. Several plausible scenarios for the abiotic emergence of membranes have been proposed (19, 20) (21). Regardless of the details of the primordial chemistry, the key feature of these proto-metabolic systems would have been simultaneous production and/or accumulation of both nucleotides and amino acids; nucleobases and simpler amino acids, at least, are readily synthesized abiogenically, under various conditions (22–26). Nucleotides, evolutionary antecedents of modern coenzymes, would function as catalysts of some of the reactions in the proto-metabolic networks, whereas other reactions could have been catalyzed by amino acids, peptides and metal clusters. Already at this stage, ATP would serve as the universal convertible energy currency. The source of energy for the primordial reproducers is a major conundrum without an unequivocal solution. The primordial membranes are unlikely to have been ion-tight as required for maintaining gradients that are converted into chemical energy in modern cells (27), so the primordial reproducers most likely were heterotrophs that made ATP by substrate-level phosphorylation.

If relatively high concentrations of nucleotides and amino acids were reached within the primordial metabolizing vesicles, synthesis of both oligonucleotides and oligopeptides at non-negligible rates could have become possible. Certain oligonucleotides can be efficient catalysts of different reactions, that is, the first, simple ribozymes. A notable case in point are self-aminoacylating mini-ribozymes which can be as small as pentanucleotides that, strikingly, catalyze self-aminoacylation almost as efficiently as modern, protein aminoacyl-tRNA synthetases (28, 29). The catalytic capacity of ribozymes is sequence-dependent, and therefore, fixation and amplification of the sequences of catalytically efficient ribozymes could become the principal driver of the evolution of replicators. Templated synthesis and ligation of oligonucleotides catalyzed by ribozymes have been demonstrated as well (30–33) although an efficient, processive ribozyme polymerase remains an outstanding goal, and additional possibilities remain open, such as involvement of peptides from the earliest stages. These processes would give rise to the first proto-replicators, most likely, relatively short polynucleotides. Even such a primitive replication process would be sufficient to kick off natural selection of proto-replicators hosted by reproducers whereby efficient catalysts would be selected along with the reproducers containing them. These ribozyme proto-replicators would become symbionts of the host reproducers. Such symbionts can be either mutualists that benefit the reproducer or parasites that exploit the reproducer. The mutualists would provide catalytic capacities that become sustainable and evolvable thanks to replication, whereas the host reproducer provides compartmentalization, resources and energy. At this stage, however, the mutualistic relationship between reproducers and replicators likely would be facultative rather than essential. The rate of RNA replication, including ribozyme-catalyzed one, is sequence-dependent, just like the rates of other ribozyme-catalyzed reactions, and hence, a different type of selection would emerge, selfish selection for the replication rate alone. Thus, parasitic replicators would inevitably evolve concomitantly with the mutualists (34, 35). These parasites would not enhance the reproduction of the host reproducers, on the contrary, decreasing their fitness through competition for limited resources, but could be problematic, if not outright impossible, to purge, in the long run.

Here we analyze an agent-based mathematical model of the co-evolution of reproducers and replicators, in an attempt to illuminate salient aspects of the evolution of the replicator-reproducer mutualism and the origin of genomes.

## Results

### An agent-based model of primordial coevolution of reproducers and replicators

#### The premises of the model

Our conceptual scenario for the origin of life as a symbiosis between reproducers and replicators is outlined in Figure 1. Reproducers are a central ingredient in this scenario. The long-term persistence of proto-metabolic networks requires some form of reproduction of the compartments encasing the reacting molecules. Division of growing lipid vesicles that strikingly resembles the division of wall-less bacteria, such as L-forms, has been demonstrated (36, 37). This is a simple, purely physico-chemical mechanism, stemming from basic physical principles, whereby vesicles become unstable after reaching a critical size and divide, not requiring complex molecular machineries that are involved in cell division in modern cells (20, 21). Although, in evolutionary biology, selection is habitually linked to replication of digital information carriers (nucleic acids), primordial reproducers, arguably, already would have been subject to a primitive form of selection. Evidently, the reproduction of the proto-metabolizing vesicles would be a far cry from the high precision process of modern cell division that is coupled to genome replication. Rather, it would be a stochastic assortment of components among the daughter vesicles (Figure 1). With this type of reproduction, random drift would necessarily play a major role in evolution, and the entire evolutionary process would resemble the stochastic corrector model that was originally proposed to describe the reproduction of primitive cells that, supposedly, contained multiple, unlinked genes (38, 39). We extend the stochastic corrector idea back to the prebiotic evolution stage that was, we surmise, the era of (pure) reproducers. Even in the evolution of such reproducers, notwithstanding the major role of drift, natural selection could set in, through the survival of the fittest vesicles, that is, the most temporally persistent ones, thanks to higher stability and/or faster growth (40).

**FIG. 1:**
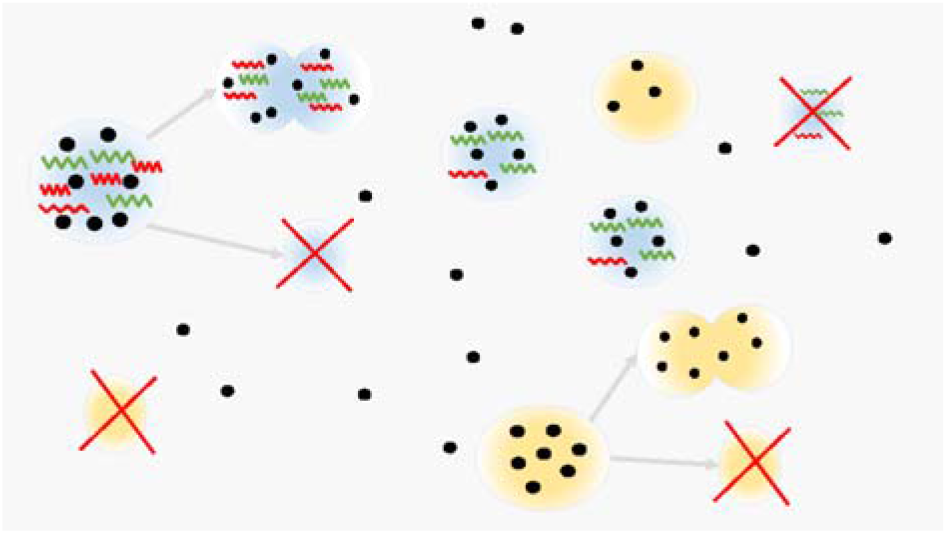
Pre-biological co-evolution of reproducers and replicators: a conceptual scenario and model framework. Protocells with (blue) and without (yellow) genetic elements (GE) compete for common resources (black circles). The GE in the protocells are either autonomous (red) or non-autonomous (green). Autonomous elements replicate themselves if the resources are present. Non-autonomous GEs replicate by interacting with autonomous elements. Both types of protocells can reproduce once their resources exceed some threshold value (in the depicted case, the arbitrarily set threshold is 7 units of resources for both protocell types). Successful reproduction yields two daughter cells. The probability of successful reproduction of the GE-containing (blue) cells depends on the composition of GE in the reproducing protocell 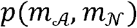, where 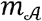 and 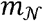 are the numbers of autonomous and non-autonomous GE, respectively. The probability of successful reproduction is constant for the GE-less (yellow) cells. If reproduction is successful, then, the daughter cells inherit the resources of the mother protocell, as well as GE, in the case of blue protocells. A reproducing cell dies in case of unsuccessful reproduction (upper left and bottom right), resulting in the dissipation of all resources and extinction of all GE. Protocells can also die due to the lack of resources (upper right blue protocell and bottom left yellow protocell).

In the symbiotic reproducer-replicator systems, competition and selection would occur at two levels: between replicators within a protocell, and between protocells carrying different complements of replicators. With respect to these two levels of selection, there would be 4 classes of replicators: 1) capable of autonomous replication and beneficial to the protocell (autonomous mutualists), 2) depending on other replicators for replication but beneficial to the protocell (non-autonomous mutualists), 3) capable of autonomous replication but useless to the protocell and incurring cost on the latter (autonomous parasites), 4) depending on other replicators for replication and useless to the protocell, thus incurring cost both on the mutualists and the protocell (non-autonomous parasites). The interactions between these distinct classes of replicators and between different replicators and protocells (reproducers) would shape the dynamics of pre-biological evolution.

#### A simple model of protocell population growth

We first consider evolutionary dynamics of cell-like reproducers (hereafter protocells) capable of resource metabolism and reproduction. There are no replicators at this stage. We assume that the protocell population is placed in a finite volume. A fixed amount of resources *R* = const is constantly supplied to that entire volume. The model is discrete, such that the time units for the protocell-level dynamics correspond to the resource supply rounds. In the given round, each protocell can acquire one unit of resources at most, thereby decreasing the total amount of resources available in the environment in the given round. The remaining resources are removed from the volume at the end of each round. Therefore, each protocell in the population is described by a resource balance (stored resources) *B^i^*, *i* = 1,2,‥ *N*, where *N* is the total number of protocells in the population. We assume that proper functionality of a protocell demands a fixed housekeeping cost Δ*E*, which is subtracted from the resource balance of the protocell at the end of the feeding phase of each round. A given protocell *i* dies if *B^i^* – Δ*E* ≤ 0 and reproduces when its resource balance exceeds a fixed threshold value *E_tr_*. Protocell reproduction is stochastic, that is, reproduction of the given protocell ends up with two cells with probability *p* (hereinafter successful reproduction probability) and no cells (mother protocell dies) with probability 1 − *p*. The resource of the mother cell is divided between the daughter cells either randomly or symmetrically (equally). In the case of random division, the resources of mother protocells are allocated randomly, with a uniform distribution, between the daughter cells. In the case of symmetric division, the resources of the mother cell are halved between the daughter cells.

The total number of protocells eventually reaches equilibrium (*N*\* denotes the population size at equilibrium) that can be approximated by considering the time variation of the total resource balance of the population of protocells (see SI Appendix A for derivation), and satisfies

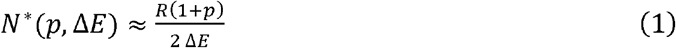

The estimate of the total population size given by (1) fails for low reproduction probabilities *p* ≤ 1/2. Indeed, in this case, each reproduction event yields less than one progeny on average, so that the total number of protocells declines over time, and thus, (1) is no longer valid. Also, (1) fails for Δ*E* ≳ 1, when the housekeeping cost approaches or exceeds the acquired resources in the given round. The estimate (1) accurately describes the total number of protocells at equilibrium for the random division case. For the symmetric division of protocells, the average number of protocells at equilibrium is slightly greater than the estimate of (1) because the average (over the population) resource balance of the protocells is greater than 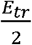 that was used to obtain (1) (SI Appendix A).

According to (1), the threshold value of the resources necessary for reproduction *E_tr_* does not affect the total population size of the protocells due to the assumption that the average resource balance of protocells is approximated by 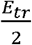. However, the threshold value explicitly enters the estimate when protocells are assumed to die (of causes other than lack of resources or reproduction failure) at each round with probability v (SI Appendix A). The threshold value plays a crucial role in the competition between populations of protocells (SI Appendix B).

We assume that any increase of the probability of successful reproduction *δp* is accompanied by an increase in the housekeeping cost *δE*. Informally, the housekeeping cost reflects the (adaptive) complexity of protocells, and we assume that more complex protocells have a higher probability of successful reproduction (41, 42). Therefore, for a given population of protocells, the change (*δp, δE*) will be evolutionarily neutral in terms of the total population size at equilibrium, that is *N*\*(*p*_0_ + *δp*, Δ*E*_0_ + *δE*) = *N*\*(*p*_0_, Δ*E*_0_) defined by (1), if the following holds for (*δp, δE*)

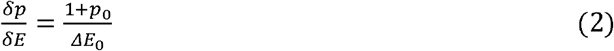

where (*p*_0_,Δ*E*_0_) are the base values of the probability of successful reproduction and housekeeping cost of the protocells, respectively. In the absence of competition, the change of the successful reproduction probability and the housekeeping cost (*p*_0_ + *δp*, Δ*E*_0_ + *δE*) would be advantageous over the initial state (*p*_0_, Δ*E*_0_) if 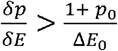, for a given population of protocells. In this case, the growth of a population is constrained only by the carrying capacity of the environment (the limited resource supply) and the housekeeping cost.

Suppose a protocell population with parameters (*p*’,Δ*E*’) competes for the common resources against a population with parameters (*p*_0_,Δ*E*_0_); the competition between protocells is indirect that is through resources only. Eq. (1) was deduced for a single population only and does not describe competition between populations. We simulated competition between two populations of protocells with different parameters and constructed the phase space plot showing the success or failure of the population of more complex protocells with a greater cost and successful reproduction probability (Fig. 2). The success in the competition is associated with their relative abundance 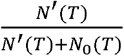 at round *T* = 3000 (one of the competing populations usually goes extinct before reaching), where and denote, respectively, the number of protocells with parameters and the pre-existing protocells with parameters respectively. We assume that the initial population sizes are. In the absence of stochastic death, in the case of symmetric protocell division, the population with a higher probability of successful reproduction and greater cost wins the competition only within a narrow range of parameters (Fig. 2a). There is a strong asymmetry between the housekeeping cost and the probability of successful reproduction: a sharp increase in the reproduction probability is required for a population to win the competition, for the given increase of the housekeeping cost. For random division of protocells, the required increase of the successful reproduction probability is smaller than in the case of symmetric division (Fig.2b). This threshold increase of the successful reproduction probability of protocells further decreases in the presence of stochastic death events for both symmetric and random division of the protocells (Fig. 2c,d). Thus, the protocells with higher successful reproduction probability and higher metabolic cost are most competitive when both the protocell division and death occur stochastically. Stochastic death favors the increase in the reproduction probability of protocells because protocells can die even if there is no lack of resources. For the given increase of metabolic cost, the increase of necessary to outcompete the base population is greater for symmetric division of protocells than for random division Fig. 2a,b). The required increase of visibly decreases even for a small death rate (same for both populations) for both symmetric and random division cases (Fig. 2c,d).

**FIG. 2:**
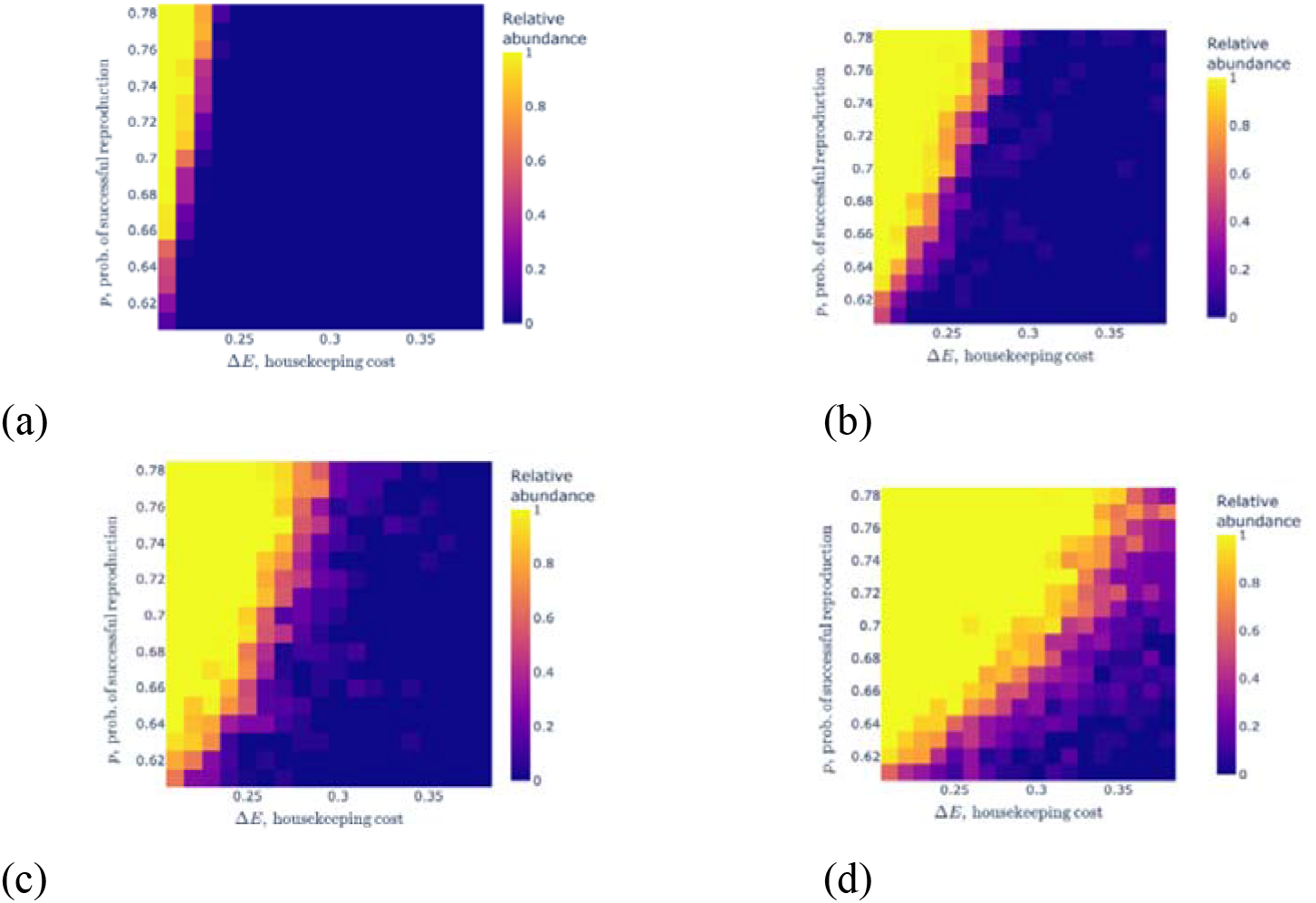
Competition of two populations of protocells with different growth parameters. Each heatmap shows the relative abundance of protocells with parameters vs the base population. The upper row panels show the outcome of the competition for symmetric division (a) and random division (b) of protocells for (no stochastic death). The bottom panels show the outcomes for symmetric division (c) and random division (d) for non-zero death rate The step sizes of the grid are. The color of each cell on the grid represents the relative abundance of the of protocells with parameters (*p*’,Δ*E*’) averaged over *M* = 20 independent simulations. The number of steps in each axis is 18. The remaining parameters are *R* = 30, *E_tr_* = 5.

To summarize the model results for the evolution of pure reproducers, we show that accounting for the metabolic cost is essential for increasing the probability of successful reproduction of protocells. Protocells with relatively low reproduction probability (and accordingly low metabolic cost) are more likely to be outcompeted by those with a slightly higher reproduction probability (and higher metabolic cots) than protocells with an already high reproduction probability. Furthermore, the increment of the reproduction probability required for a protocell population to win over a pre-existing population of protocells is larger for symmetric division than for random division of protocells, for the given increment of housekeeping cost. Both random division of protocells and stochastic death shift the trade-off between the advantages of higher successful reproduction probability and the increasing metabolic cost towards the former. These observations on pure reproducers are directly relevant for the origin of the mutualistic symbiosis between protocells and GE. The presence of GE in the protocells can have a crucial impact on the metabolism (10, 43). Below, for simplicity, we assume that the positive feedback of the GE is manifested by the increased probability of successful reproduction of the host protocell, without explicitly considering metabolism. Indeed, it is the trade-off between the successful reproduction probability and the metabolic cost that enables advantageous feedback mechanisms between the protocells and GE, whereby GE improve the error-prone reproduction process of the primordial reproducers (protocells), whereas the protocells provide resources for the replication of GE, thus, increasing the housekeeping cost, as discussed in the subsequent sections.

#### Evolution of genetic elements in the absence of (proto)cells

Before addressing the coevolution of genetic elements with protocells, we consider the idealized (even if biologically unrealistic) case of evolution of GE in the absence of protocells (compartments). In the model, there are two types of GE, autonomous and non-autonomous replicators. The autonomous GE can replicate themselves as well as the non-autonomous GE, whereas the non-autonomous GE can replicate only through interaction with the autonomous ones and do not contribute to the replication of the latter. The replication of both types of GE is assumed to be possible only in the presence of the necessary resources, and along with replication, GE can also die. Although autonomous GE can mutate into non-autonomous ones, we consider such mutations to be rare and, for simplicity, do not include these in the model.

The GE are placed in a well-mixed volume and directly consume the resources which are supplied at a fixed rate. The following elementary processes describe the minimal model of resource supply and birth and death of GE (44):

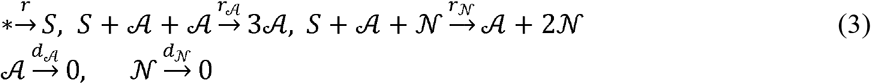

Here, *S* denotes the resources (substrates) available in the environment, 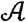 and 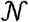 denote autonomous and non-autonomous replicators, respectively. 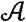 serves as a replicase for the replication of both its own type and non-autonomous type of GE. Because a molecule cannot simultaneously function as both the template for replication and the replicase, the reaction for the replication of autonomous GE includes two copies of 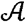, one for each of these functions, whereas the reaction for non-autonomous GE requires only one 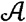, the replicase and one 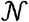, the template. From (3), we can obtain the deterministic description of the evolution of the number of GE and the amount of resources (see SI Appendix C). The dynamical system obtained by (3) is a consumer-resource model (45–47) that can have three distinct equilibria. In the first equilibrium, 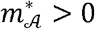 and 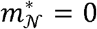 (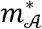 and 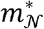 are the numbers of autonomous and non-autonomous GE at equilibrium), that is, only autonomous GE are present in the environment. The second equilibrium is coexistence of both types of GE, 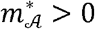 and 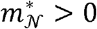. The third equilibrium corresponds to the extinction of both types of GE, 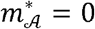 and 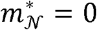. The extinction of both types occurs under the condition

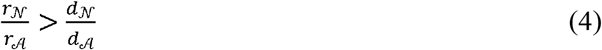

The coexistence of both types is possible under the fine-tuned condition, that is, when (4) holds as equality, and the autonomous-only equilibrium can be reached if (4) is reversed (see SI Appendix C).

The extinction of both types of GE is due to the overtake of non-autonomous replicators in the environment, suppressing the replication of autonomous GE. The autonomous replicators go extinct due to the death events, eventually resulting in the collapse of the population of non-autonomous replicators as well. Thus, both types of GE die out and the entire population of replicators collapses (4). This model describes the birth-death process of GE in a spatially homogenous environment. Spatial heterogeneity in the environment can result in compartmentalization of GE, allowing the coexistence of both types (43,70). Below, we assume 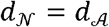, and consider two cases: 1) the reaction rates are almost equal 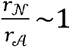 (but (4) still holds) and 2) the extreme case 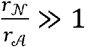.

#### Competition between protocells containing and lacking genetic elements

In the previous sections, we separately described the competition among protocells lacking GE (pure reproducers) and the competition among GE in the absence of protocells (pure replicators), where resources are supplied directly. We now explore the case most relevant for the origin of life, when the interactions among GE, described by (3), occur inside protocells and there is feedback between protocell reproduction and GE replication. We assume that GE can survive only in the protocells, hence the death of a protocell leads to the death of all its GE as well. The GE use the resources of the host protocell for the replication, and the replication of any GE is associated with an additional cost *E_c_* « 1 (much smaller than the acquired resources per round) which is subtracted from the protocell’s resource balance. Therefore, intracellular replication of GE increases the housekeeping cost for the host protocells. Obviously, under these conditions, protocells that harbor GE will lose the competition against those that lack GE unless at least some of the GE are beneficial to the protocells. We therefore assume that the presence of mutualist GE in a protocell increases the probability of successful reproduction whereas parasitic GE only incur cost. The interplay between the opposing effects of GE on the reproduction of protocells defines the evolutionary outcome for the entire system. In this section, we analyze the interaction and evolution of only two of the four classes of GE defined above, Class 1 (autonomous mutualists) and Class 4 (non-autonomous parasites) GE. The case of non-autonomous mutualists and autonomous parasites is addressed in the next section.

The initial number of GE in protocells is given by the Poisson distribution with parameter μ

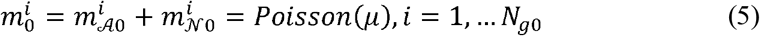

where *N*_*g*0_ is the initial number of protocells containing GE (at the start of the first round of resource supply). The initial number of mutualists in each protocell is defined by a randomly (with uniform distribution) chosen integer from 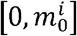, that is the number of GE in each protocell is the same for the given simulation, defined by (5), only the fraction of mutualists varies in protocells. We assume that there is a time-scale difference between the protocell reproduction and competition, on the one hand, and the intracellular birth-death process (replication) of the GE, on the other hand, such that the latter process is much faster than the former one. The protocell-level competition is governed by the rounds of resource supply to the environment. Let us denote by *K* the number of GE replication and death events (3) that occur in a protocell in each round of resource supply. In a given round, the GE can use *KE_c_* resources of the protocell at most (when no death but only replication of the GE occurs). In the extreme case *K* → ∞, the intracellular replication of the GE goes to steady state in each round, resulting in either a GE-less protocell or the death of the protocell together with its GE due to the resource exhaustion.

Here we assume that the protocells benefit from the mutualist GE they contain through an increased probability of successful reproduction. The probability of successful reproduction of a GE-containing protocell linearly depends on the fraction of mutualists it contains

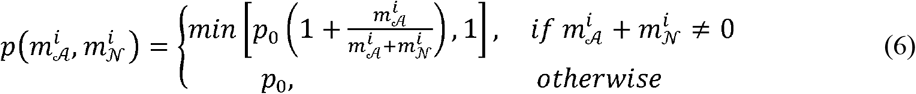

Here *p*_0_ is the probability of successful reproduction of the protocell in the absence of GE. The probability of successful reproduction is the same for the protocells containing only parasites 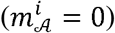 and the GE-less protocells 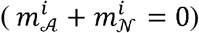, but if both mutualist GE and parasites are present in the protocell (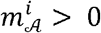 and 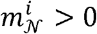), then, the increase of the number of parasites offsets the benefits of mutualist GE (6). Thus, proliferation of parasites impacts the protocell in two ways: they consume resources during replication (which occurs only in the presence of mutualists) and also decrease the probability of successful reproduction of the protocell decreasing the fraction of mutualists among the GE (6).

We consider three reproduction scenarios that differ with respect to the allocation of the resources and GE of the mother cell between the daughter cells.

1. *Random allocation of GE and proportional allocation of resources*. In this scenario, the number of mutualists and parasites in one of the daughter protocells is randomly (with discrete uniform distribution per type of element) drawn from the dividing mother cell. The resources are distributed proportionally to the total numbers of GE in daughter protocells

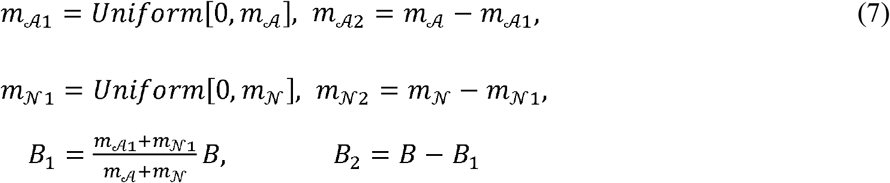 The daughter protocells are denoted by 1 and 2, 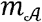 and 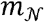. are the total number of autonomous and non-autonomous GE in the mother protocell at the time of division, *B* is the resource balance of the mother cell at the division. As shown previously by other, this random division mechanism (13) is the most favorable among several other mechanisms of binary division, with respect to the appearance of mutualists-only protocells (48).
2. *Random allocation of resources and binomial distribution of GE*. Here the amount of resources in the daughter cells is determined by the random division of mother cell with uniform distribution. The numbers of GE in each of the daughter protocell are then determined by the binomial distribution with the parameter defined by the distribution of the resources between the daughter cells. In more concrete terms, such allocation of both the resources and the GE could be linked to the distribution of the relative sizes of the daughter protocells.

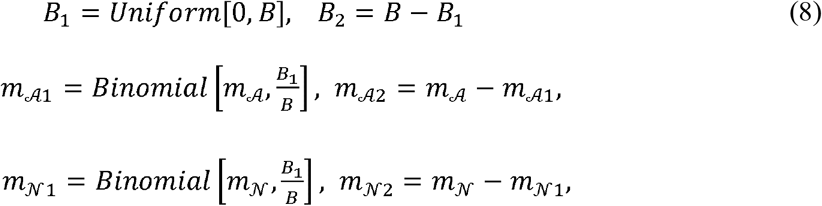
3. *Symmetric distribution of both GE and resources*. Here both the GE and resources of the mother cell is (almost) equally divided between the daughter cells.

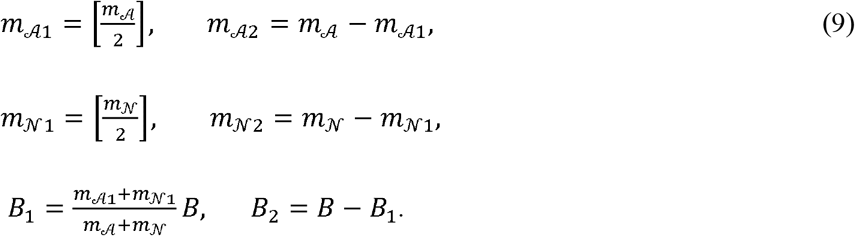

where 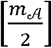 is the integer part of the ratio 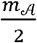. Symmetric division is considered here as an extreme case, even if not necessarily realistic one at the protocell stage of evolution.

Consider competition of GE-containing protocells against GE-less protocells under the random division scenario (7) (Figure 1). If a GE-containing protocell loses those elements, it joins the population of GE-less protocells. The protocells that only contain parasites eventually lose them due to the deaths of GE and impossibility of parasite replication in the absence of mutualists. The basal housekeeping costs (in the absence of GE) and threshold values are the same for both type of cells.

Random allocation of GE among the progeny and death of GE eventually will result in the appearance of some protocells that carry only mutualists. The appearance of these mutualist-only protocells depends on the reproduction threshold value *E_tr_* (time interval between two successive divisions of the protocells depends on the *E_tr_*) and the average number of intracellular elementary processes *K* in the given round of resource supply. Both quantities control the balance between mutualists and parasites. Indeed, in the extreme case 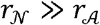, it can be assumed that only birth of parasite occurs, hence the increase of the number of parasites will be proportional to the *KE_tr_*. Also, increase of *K* causes a concomitant increase of the housekeeping cost of the protocell by approximately *KE_c_*.

The fractions of autonomous mutualists and non-autonomous parasites in the protocell population at round *t* are

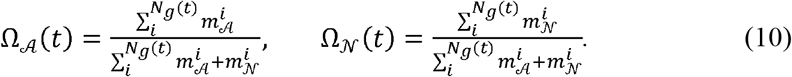

All other parameters being equal, the outcome of the competition critically depends on *K*. For *K* = 1, that is, when replication of GE is coupled with the protocell reproduction, GE-containing protocells win the competition (Fig. 3a), and following a brief initial surge of parasites, mutualist-only protocells take over (Fig. 3b). In contrast, at *K* = 15, parasites take over in the initial phase of the competition, effectively precluding the appearance of mutualist-only cells. As a result, GE-less protocells win the competition (Fig. 3c) whereas all GE die off (Fig. 3d).

**FIG. 3:**
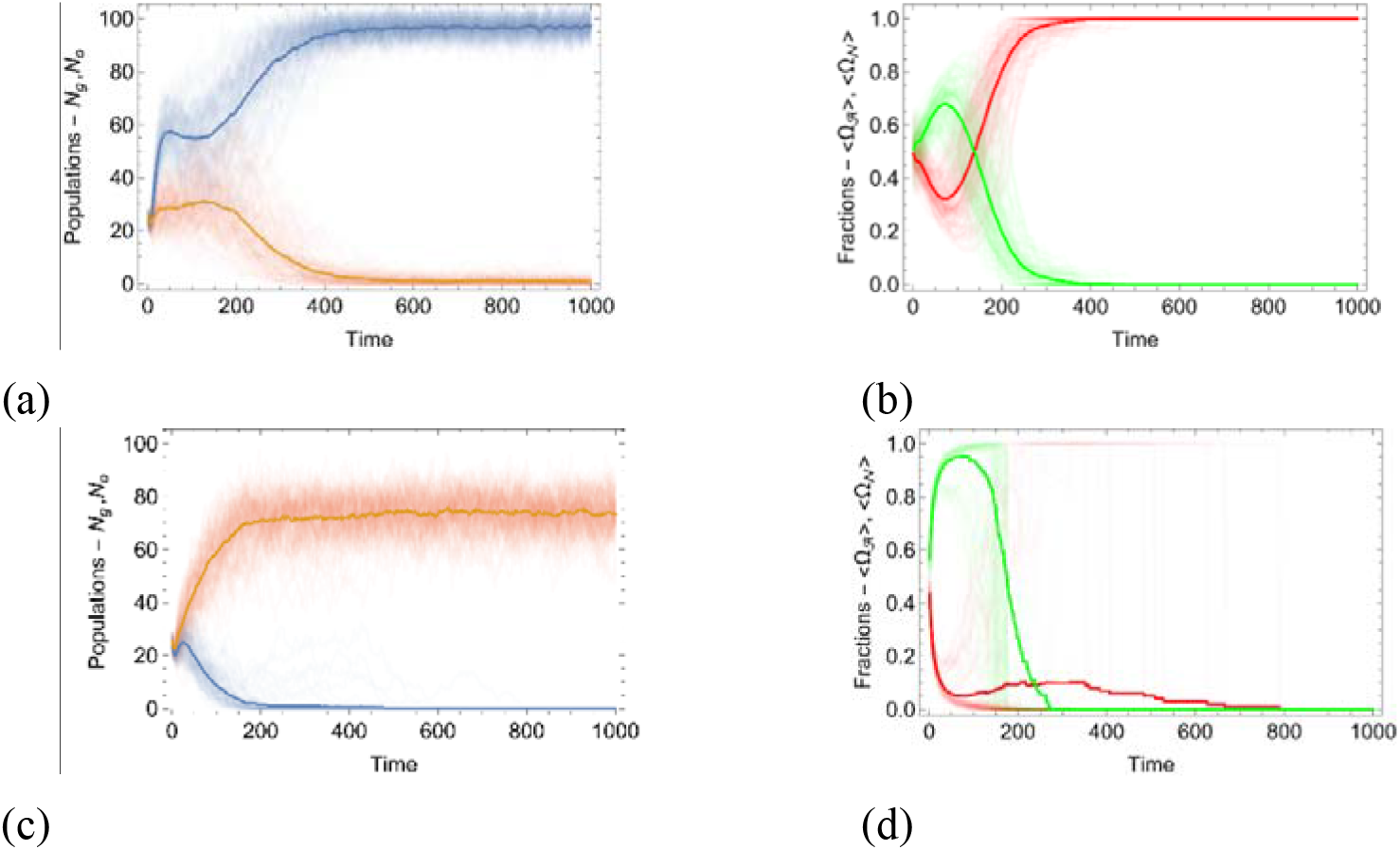
Competition of protocells containing genetic elements against protocells lacking genetic elements under the random division scenario. The simulations were performed under the random division scenario (7) and assuming 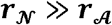. The plots on the left show the time dependency of the total number of the protocells with (blue lines) and without (orange lines) GE, denoted by *N_g_* and *N_o_* respectively, for (a) *K* = 1 and (c) *K* = 15. The thick lines show the size of the populations averaged over an ensemble of 100 realizations. The plots on the right show the time dependency of the fraction of autonomous mutualists (red lines) and non-autonomous parasites (green lines) for (b) *K* = 1 and (d) *K* = 15. The thick lines show the behavior of fractions of GE (10) averaged over the ensemble. The remaining parameters of the model are as follows: 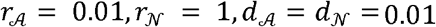, *R* = 30, Δ*E* = 0.3, *E*_c_ = 0.01, *p*_0_ = 0.6, μ = 100, *E_tr_* = 5.

The probability of successful reproduction (6) is greater in the initial phase of the dynamics due to the random distribution of the mutualists among the protocells. On average, the initial fractions of mutualists and parasites are almost equal in the population (Figs. 3c,d). The population of GE-containing protocells initially grows faster than the population of GE-less protocells, regardless of the eventual outcome (Figs. 3a,c), and similarly, the fraction of parasites initially grows in all cases (Figs. 3b,d). By contrast, at low *K* values, the mutualist-only protocells emerge stochastically, due to the random protocell division, and eventually win the competition due to the selective advantage conferred by the mutualists (6).

The values of *K* and *E_tr_* determine the outcome of the competition for other reproduction scenarios as well. Figure 4 shows the results of the competition between GE-containing and GE-less protocells depending on the values of *E_tr_* and *K* for all three scenarios, described by (7), (8) and (9). Here, we took snapshots of the number of GE-containing protocells and GE-less protocells at sufficiently large time *T*. For each simulation, the relative abundance of GE-containing protocells 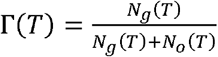 was calculated, and then averaged over the ensemble of 100 independent simulations, that is, independent realizations of the population dynamics

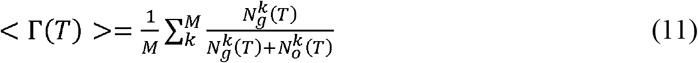

**FIG. 4:**
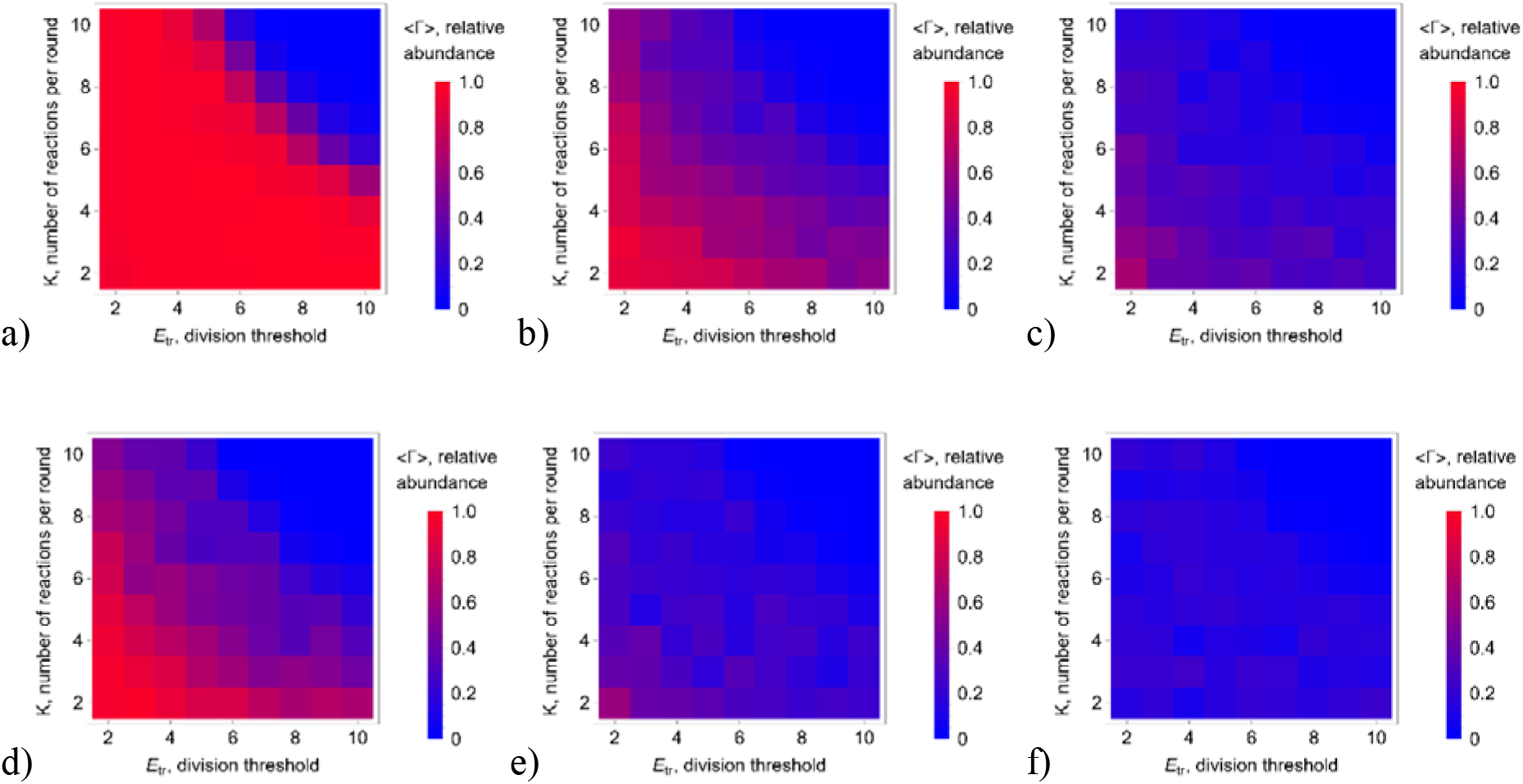
Dependency of the relative abundance of GE-containing protocells on *K* and *E_tr_* for the three division models. From left to right, the panels correspond to the random (7) (a,d), binomial (8) (b,e) and symmetric (9) (c,f) division scenarios, respectively. The rows (a, b, c) and (d, e, f) show the outcome of the competition for 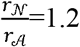 and 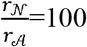, respectively, where 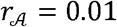. The bottom left corner on each subfigure corresponds to (*K*, *E_tr_*) = (2,2), and the step of the heatmap is 1. The snapshots are taken at *T* = 3 · 10^3^ and averaged over *M* = 10^2^ simulations. The remaining parameters are the same as in Fig.3.

The outcome of the competition is presented for both 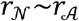(such that (4) holds) (Fig.4 a,b,c) and for 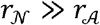 (Fig. 4 d,e,f). The GE-containing protocells can win the competition when reproduction and replication are on the same time-scale, that is, at small values of both *K* and *E_tr_*. Random division (7) (Fig.4a,d) is the most advantageous scenario for the GE-containing protocells for both 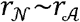 and 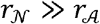. Even in the extreme case 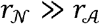, (7) allows winning of the GE-containing protocells for a large region of the phase space (Fig.4d). By contrast, for binomial division (8), GE-containing protocells can win only within a small region of the phase space (Fig.4e) corresponding to the smaller values of (*K*,*E_tr_*), and for symmetrical division (9), GE-containing protocells always lose (Fig. 4f).

The success of GE-containing protocells in the competition for common resources is due to the appearance of mutualist-only protocells in the population. The probability of the appearance of mutualist-only protocells is greater in the case of random division (7), than for the binomial (8) or symmetric (9) division. Indeed, for scenario (7), the probability that the given daughter protocell will contain mutualist-only GE (at least two such elements 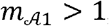, which is the minimum requirement for replication under (3)) is

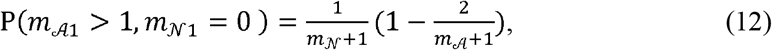

where 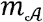 and 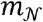 are the numbers of autonomous and non-autonomous GE in the mother protocell at the time of reproduction, respectively. For the random allocation of resources and binomial distribution of GE (8), the corresponding probability is (see SI Appendix D for details)

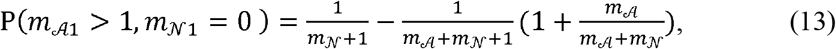

The value given by (13) is smaller than that given by (12) as long as 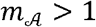 and 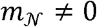. In the case of symmetric division (9), the appearance of mutualist-only protocells due to the reproduction of the mother protocell is impossible for 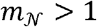. Mutualist-only protocells can still emerge even in this scenario as a result of stochastic death of parasites or due to the initial distribution of GE in the protocells (5). The initial distribution of GE in the protocells impacts the appearance of mutualist-only protocells. The smaller the initial number of GE in protocells, defined by parameter *μ* in (5), the more likely is the appearance of mutualist-only protocells (see SI, Appendix E).

#### Non-autonomous mutualists and autonomous parasites

We further examined a distinct version of the model in which protocells contained non-autonomous mutualists (Class 2 replicators) and autonomous parasites (Class 4). In this case, the probability of successful reproduction of the protocells (6) increases with the fraction of (non-autonomous) mutualists (swapping 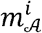 and 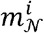 in (6)). Both protocell-level and GE-level selection favor the appearance of mutualist-only protocells due to (6) and (4), respectively. However, in this case, because the mutualists are incapable of replication, such protocells will lose all GE (3). The appearance of protocells containing only autonomous parasites is unfavorable as well because these GE provide no advantage to the protocell and, on the contrary, being able to replicate, incur a cost by consuming resources. In this case, to enjoy a sustainable competitive advantage, the GE-containing protocells have to carry both types of GE. The probability of successful reproduction (6) of such protocells will be lower than that of protocells containing only non-autonomous mutualists, and so they will be outcompeted, irrespective of the division scenario (Fig. 5).

**FIG. 5:**
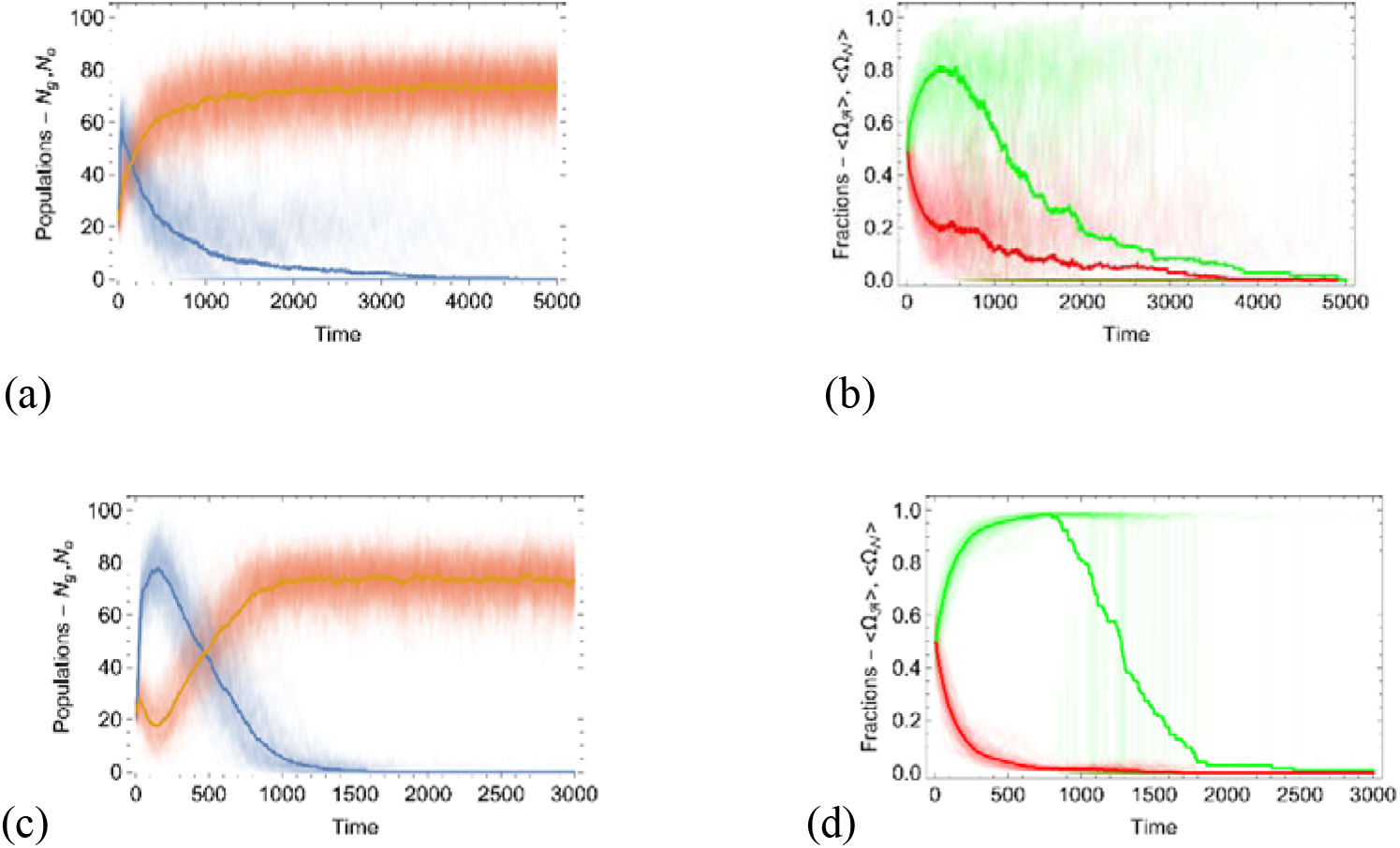
Competition of protocells containing genetic elements against protocells lacking genetic elements: non-autonomous mutualists and autonomous parasites. Time dependency of the total number of the protocells with (blue lines) and without (yellow lines) GE, denoted by *N_g_* and *N_o_* respectively, for the division scenarios (7) and (9), (a and c, respectively). The thick lines show the size of the populations averaged over an ensemble of 100 realizations. Time dependency of the fraction of autonomous parasites (red lines) and non-autonomous mutualists (green lines) for (b) and (d). The thick lines show the behavior of fractions of genetic elements (10) averaged over the ensemble. The model parameters are as follows: *E_tr_* = 5, *K* = 1, 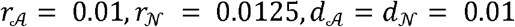, *R* = 30, Δ*E* = 0.3, *E_c_* = 0.01, *p*_0_ = 0.6, μ = 100, *E_tr_* = 5.

The survival of non-autonomous GE is impossible under the assumption that the protocells divide rarely (large division threshold *E_tr_*), mimicking (3) in the absence of protocells. The same conclusion holds also for the case when the non-autonomous mutualists are at a disadvantage in the GE-level selection, that is, when (4) is reversed. In this case, the autonomous parasite will outcompete the non-autonomous mutualists in the protocells, decreasing the possible benefit to protocells due to (6). Even if mutualist-only protocells appear, they will lose their GE due to the lack of autonomous parasites. Conversely, protocells containing only autonomous parasites will lose the competition against GE-less protocells. Thus, with this combination of GE types, there is no path for GE-containing protocells to take over the population. Extrapolating these findings for the case of protocells containing all 4 types of GE, takeover by GE-containing protocells is only possible through the emergence of protocells containing only autonomous mutualists, regardless of the initial combination of GE types.

#### Synopsis of the model results

We now briefly summarize the results of our modeling of the coevolution of reproducers and replicators. We focus on the cost-benefit analysis of feedback mechanisms between protocells and GE. First, we developed a model that describes the population dynamics of protocells capable of resource metabolism and reproduction, in the absence of any GE. We considered the role of various parameters (threshold of the resource amount necessary for reproduction, housekeeping cost, probability of successful reproduction, symmetric vs. random reproduction of protocells) in the protocell (reproducer) level dynamics in the absence of any GE. Then, we addressed the case of evolution of two types of GE, autonomous and non-autonomous, in the absence of protocells. In the model, the replication rate of non-autonomous genetic elements is greater than that of autonomous elements. We focus, in particular, on the part of the parameter space where non-autonomous GEs outcompete autonomous GE, leading to the extinction of both types in the absence of protocells. Then, feedback is introduced between protocells and GE whereby the GE interact among themselves within a protocell (including both competition and autonomous GE providing the replication machinery to non-autonomous GE) and also with the protocell. The GE use the resources of the host protocell for their replication, thus incurring an additional cost on the protocell. However, the presence of mutualists in the protocell increases the probability of successful reproduction. The main focus of this work is the competition between GE-containing and GE-less protocells. We observed that selection favors GE-containing protocells when the protocell reproduction and GE replication occur on the same time scale, best of all, are precisely coupled, and when the resource threshold for reproduction is low. Random division of resources and GE is advantageous compared to symmetrical division for GE-containing protocells because randomness provides for the appearance of mutualist-only protocells that take over the population. We further considered the case of non-autonomous mutualists and autonomous parasites where replication-competent, autonomous GE carry no benefit for the protocell whereas the non-autonomous GE do. In this case, the GE-containing protocells lose the competition, since the non-autonomous mutualists are not able to replicate in the absence of autonomous parasites.

## Discussion

Origin of life, or more precisely, the origin of cells, the universal evolving units of life, remains a fundamental enigma (49). The information transmission pathways of modern cells that underpin evolution are exquisitely complex and themselves must have evolved under selection (50, 51). Motivated by these considerations and by the fundamental distinction between the two types of evolving biological entities, reproducers and replicators (4, 39), we modeled here the origin of cells as a mutualistic symbiosis between protocellular reproducers and primordial replicators. Protocells are commonly perceived as entities that already contained replicating RNA molecules within lipid membrane-bounded vesicles (52). We argue, however, that a stage preceding the origin of cells and involving selection for persistence (40) among pure reproducers devoid of any GE is inescapable in prebiological evolution. In our scenario, these protocellular reproducers become incubators for primordial replicators (GE) (29, 53) that increased the fitness of the protocells carrying such replicators, enhancing protocell reproduction, and only subsequently, assumed coding functions.

Replicators, however, present an inherent, major problem in that their evolution inevitably gives rise to parasitic elements that hijack the replications machinery of autonomous elements (34, 35). In homogeneous, well mixed systems, the parasites that replicate faster than autonomous elements tend to take over, leading to the collapse of the entire replicator ensemble (44). Compartmentalization can substantially change the evolutionary dynamics of autonomous elements and parasites, preventing the takeover by parasites and stabilizing the system (44, 54-57). Furthermore, previous modeling studies strongly suggest that replicators are more likely to survive within protocells compared to surface-based spatial systems (17).

Here, we explored a mathematical model of the evolution of replicators (GE) within protocellular reproducers seeking to define the conditions that favor the selection of protocells carrying GE and eventual emergence of *`bona fide`* genomes. Under the assumption that mutualist GE conferred selective benefits onto the host reproducers, within which such GE replicated, we identified three key conditions for the fixation of the genetic system in evolution. First, the GE level dynamics (replication and death) has to be coupled to the reproduction of the protocells. Counterintuitive as that might seem, GE replication at rates substantially higher than the reproduction rate of the protocells leads to the extinction of the GE. Informally, this requirement stems from the need to avoid exhaustion of the resources available for protocell reproduction by uncontrolled replication of the GE. Second, along similar lines, the threshold amount of resources required for reproduction has to be sufficiently low for the protocells to afford GE replication. Third, the distribution of the GE between daughter cells has to be random rather than symmetrical, to enable the emergence of mutualist-only protocells. A similar conclusion has been previously reached for a model of group selection of replicators within cells (58). Importantly, it should be noted that the model is contingent on the mutualist GE not being essential for protocell reproduction which appears to be an appropriate condition for the primordial stage of evolution.

We further investigated a version of the model, in which only parasites but not mutualists were endowed with the replication capacity. In this case, obviously, a reproducer-GE system could potentially persist and be competitive only if it contained both parasites and mutualists. The probability of successful reproduction of such protocells will be lower than that of protocells containing only non-autonomous mutualists, and so they would be outcompeted by the latter. However, protocells containing only non-autonomous GE that cannot replicate will, evidently, lose their GE content. Thus, this combination of GE types is not conducive to the selection of GE-containing protocells.

These simple, intuitive constraints on the early steps in the evolution of GE have substantial implications for the origin of genomes and modern-type cells. The first of these pertains to the origin of large genomes, on the scale of the genomes of the extant bacteria and archaea. There is little doubt that the first GE were small, on the order of a kilobase, at most. Our model shows that GE-containing protocells could stand a chance in the competition with GE-less ones only when all or at least a large fraction of the GE encoded their own replication machinery. Thus, at the early stages of the genetic system evolution, the key role apparently belonged to GE resembling modern RNA viruses, especially, those that do not encode any structural proteins, but only the enzyme required for replication, such as narnaviruses or mitoviruses (59, 60). Protocells that harbored ensembles of mutualists would encompass multiple versions of the replication machinery. This excess of sequences dedicated to replication would engender selective pressure for joining genetic elements and eliminating the redundancy, saving resources and facilitation coordination of replication with protocell division. The second major corollary is the requirement for coupling the replication of mutualist GE to the protocell division. Such coupling is a universal feature of modern cells, and our findings strongly suggest that it was selected for from the earliest stages of (pre)cellular evolution. The third implication concerns the origin of dedicated defense systems against parasites. Defense systems are extremely abundant and diverse in modern prokaryotes where they account for a considerable, probably, still underestimated fraction of the genome. (61, 62). At the primordial stages of evolution, when protocells divided stochastically, the mutualist-only protocells would win the competition against those lacking GE or those infested by parasites. However, once the fusion of primordial mutualist replicators gave rise to large genomes present in a single or few copies per cell, symmetrical division would evolve driven by the selection for accurate segregation of genomes into the daughter (proto)cells. However, the problem with symmetrical division is that it makes (proto)cells vulnerable to parasite onslaught as parasites would persist after invading or evolving from mutualists by mutation. Therefore, it appears that defense mechanisms, conceivably, those based on specific recognition of parasite sequences, would coevolve with symmetrical cell division mechanisms, being a pre-requisite for the long-term survival and evolution of cells.

The present scenario for the origin of cells is not inconsistent with the RNA world hypothesis (52, 53, 63). We stress, however, that the primordial RNA world must have evolved within pre-existing, metabolically active, membrane-bounded protocells (reproducers) as proposed previously by Copley, Smith and Morowitz (64).

Even if quite general, the model of the origin of life presented here suggests many avenues for experimental testing. In particular, experimental modeling of the origin of replicators within reproducers, that is, membrane vesicles encompassing proto-metabolic networks producing nucleotides and amino acids, and potentially, oligonucleotides and peptides, might not be far beyond the capability of modern laboratories (65, 66).

### Brief methods

For the protocell dynamics, the steps of the simulations are as follows. In the absence of GE, the protocell is described by the tuple of resource balance of the given cell and the type of the cell {*B^i^*, *type*} that defines the model parameters for that protocell (Δ*E, p, E_tr_* and so on). GE-containing protocells are described by {*B^i^*, *type*, 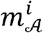, 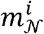}. Thus, the whole population of protocells is described by the list of tuples. At the beginning of each round, the orderings of the tuples are randomly permuted to ensure swell mixed condition. A constant amount of resources *R* is supplied in the environment. The resource balance of the *i*-th cell of the population will be either 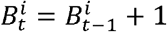 if there are still available resources in the environment before feeding, or will remain the same 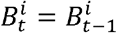 if the resources are already exhausted at the given round. After feeding, each protocell in the population pays the housekeeping cost defined by its type. If, after paying this cost, the resource balance of the protocell is not positive 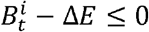, then the protocell is removed from the population.

The reproduction phase of the protocell starts when the resource balance is greater than the threshold value for reproduction defined by the type of the protocell, that is 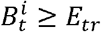. The reproduction of the protocell ends up in two progeny protocells with probability *p* and no progeny with 1 − *p* (note, that for GE containing cells 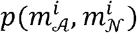). That is, if the randomly generated number in the unit interval is less than *p*, then, new element is added in the list with the same type, and the resources of the reproducing protocell is defined according to the reproduction scenario. Then, the number of tuples of the given *type* is selected from the main list representing the number of protocells of the given *type* in the end of each round.

The intracellular dynamics of GE, that is, the birth and death process described by (3), is modeled using the Gillespie method (66), excluding the time of the occurrence of birth-death elementary processes because it is assumed that the number of elementary processes in each GE-containing protocell is equal to *K* at any given round of resource supply. The propensities of the elementary processes are constructed first. For the birth of autonomous GE, 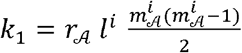, where 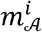 is the number of autonomous GE in the protocell, and 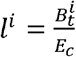 describes the possible number of GE that can be made from the resources of the cell (*l^i^* = 0, if 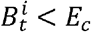). A birth of a autonomous element results to the following changes 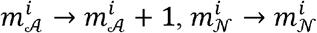 and *l^i^* → *l^i^* − 1. Similarly, for the birth of non-autonomous elements, the propensity is 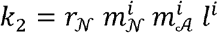. The amount of resources and the numbers of GE and resources change according to 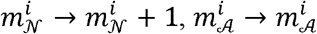 and *l^i^* → *l^i^* − 1. The propensities of death processes are 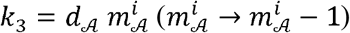 and 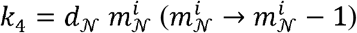 for autonomous and non-autonomous elements, respectively. Then, a random number is generated from the *∊* ∈ [0,1] interval, and a reaction is chosen for which the following condition holds 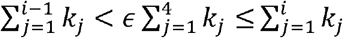, where *i* = 1,‥4. Then, the number of GE in the protocell is updated according to the chosen process. The same steps are repeated *K* times in each round of the resource update in the environment. Note that the resource balance in the protocells is governed by the protocell level dynamics described above.

All other aspects of the simulations are presented in the Results section.

## Acknowledgements

S.B., Y.I.W. and E.V.K. are supported by the Intramural Research Program of the National Institutes of Health of the USA (National Library of Medicine). A.A. and A.K. are supported by State Committee of Science of Armenia, grant No. 21AG-1C038. P.L-G is supported by basic funding of the French Centre National de la Recherche Scientifique.

