## Supporting information for "Coevolution of reproducers and replicators at the origin of life and the conditions for the origin of genomes"

***Appendix A.***

***Coarse grained model for the evolution of a single protocell population***

The resource balance of the whole population in any given round is $B_{tot}=\sum_{i=1}^{N} B^{i}$, where $N$ is the number of protocells in that round. Let us assume that there is no difference in the resource balances among the protocells, such that $B_{tot}=NB_{av}$. For successful reproduction probabilities $p >½$, the change of the total resource balance of the protocell population in round $t$ is

$N{\Delta B}_{av}(t) = R- N\Delta E- \lambda(1- p)N B_{av}$*.*	(A1)

The $R$ term in (A1) is the resource intake by the population (for large $N$, we can assume that the population consumes all available resources $R$ in that round). The second term $N\Delta E$ is the total housekeeping cost. The last term is dissipation of resources caused by unsuccessful reproduction (death) of protocells. Here we assumed that for large $t$, the reproduction of cells is a Poisson process with the mean $\lambda=\frac{R/N}{E_{tr}}$ –, where $1/\lambda$ is the mean number of rounds required to exceed the threshold value. Note that population growth decrease $\lambda$, that is the weight of the resource dissipation by failure in the reproduction events become decreases. For *t* → ∞, the right side of (A1) is equals zero, that is, the total input and output of the resources are balanced. We approximate $B_{av}(\infty)\approx\frac{E_{tr}}{2}$ assuming that feeding and reproduction of cells are uncorrelated; this approximation works best for high reproduction probability $p$.

The distribution of resource balance values in the population is more narrowly concentrated around $\frac{E_{tr}}{2}$ in the case of symmetric division of the resources between the daughter cells than in the case of random division of the resources. For the symmetric case the average resource balance of the cells is $B_{av}\left( \infty\right)\geq\frac{E_{tr}}{2}$ , since at any given round *t* → ∞ the population of protocells consist of the newborn progenies that have been born in the previous round and the cells that have not been reproduced. The resources of newborn progenies is larger than $\frac{E_{tr}}{2}$. Hence, the average resource balance will be larger for the symmetric division, consequently the average number of protocells will be greater than in the random division case (A1). From (A1), we derive the total number of cells $N^{*}$ for $t\to\infty$

$N^{*}=\frac{R(1+p)}{2 \Delta E}$	(A2)

In Fig.S1, the time dependence of the average number of protocells is illustrated for different values of model parameters. The estimate of the total population size given by (A2) fails for small reproduction probabilities (Fig.S1b) and also for $\Delta E\approx1$, when the housekeeping cost is almost equal to the acquired resources in the given round. In (A2), the number of cells at equilibrium does not explicitly depend on the threshold value of the resources necessary for reproduction of the cells. Therefore, we observed a substantial discrepancy between the average total number of cells and the estimate given by (A2) (see Fig.S1c). Indeed, for the smaller threshold values, the assumption of reproduction being a Poisson process fails although the average of the resource balances of the protocells remains close to $\frac{E_{tr}}{2}$ . However, for larger $E_{tr}$*_,_* the discrepancy decreases, and (A2) can be used as an estimate of the average population size.

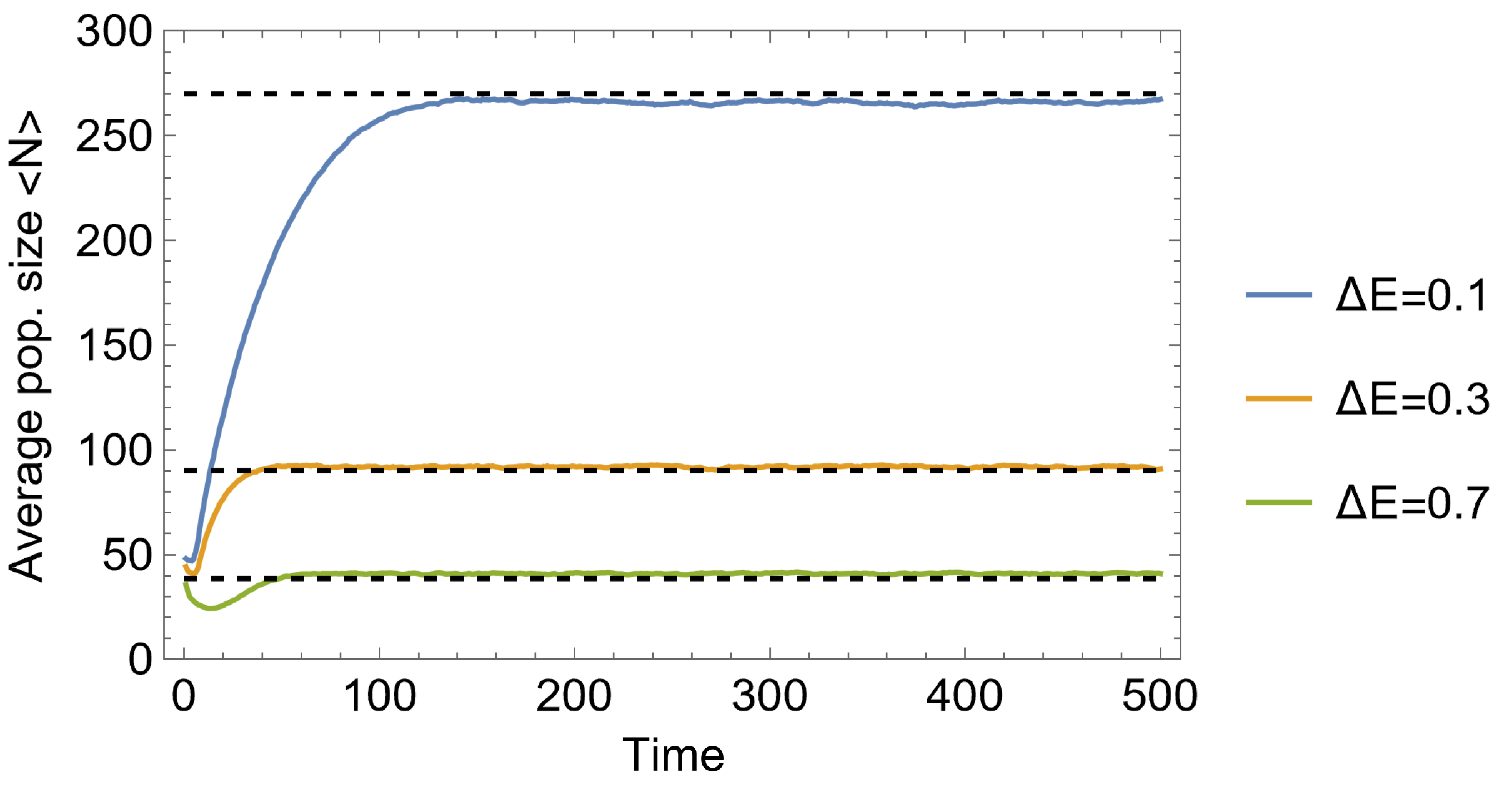

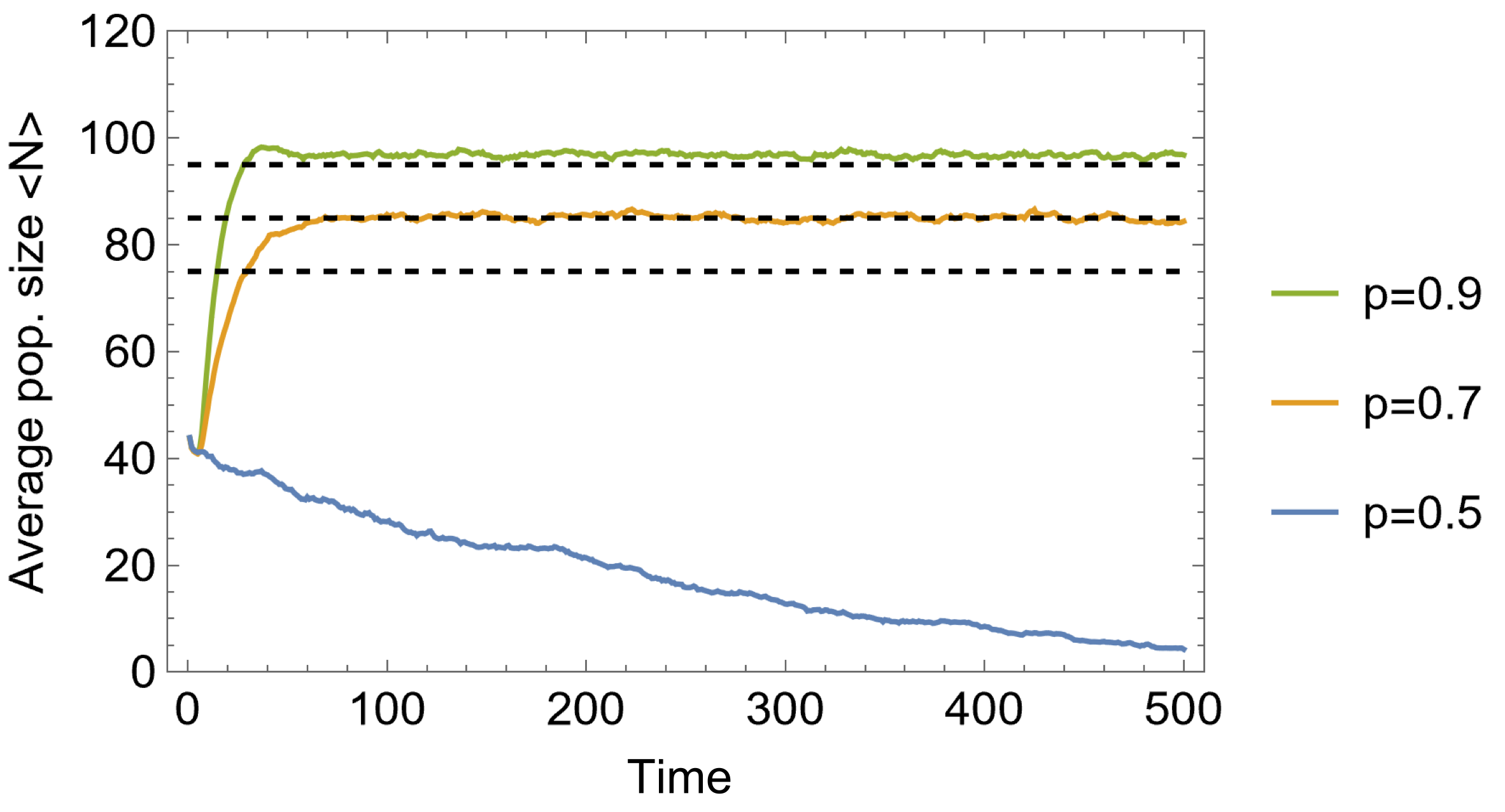

(a)	(b)

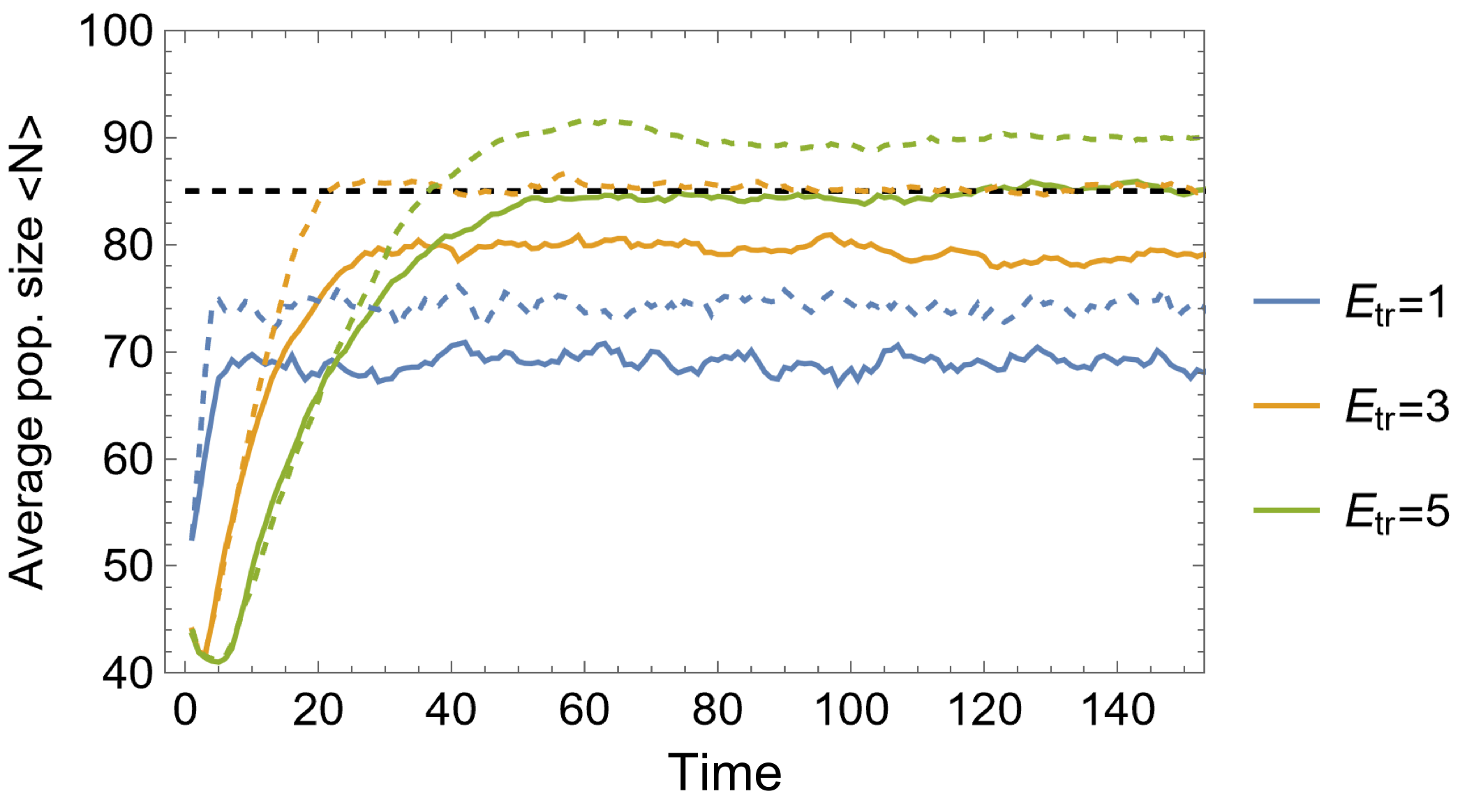

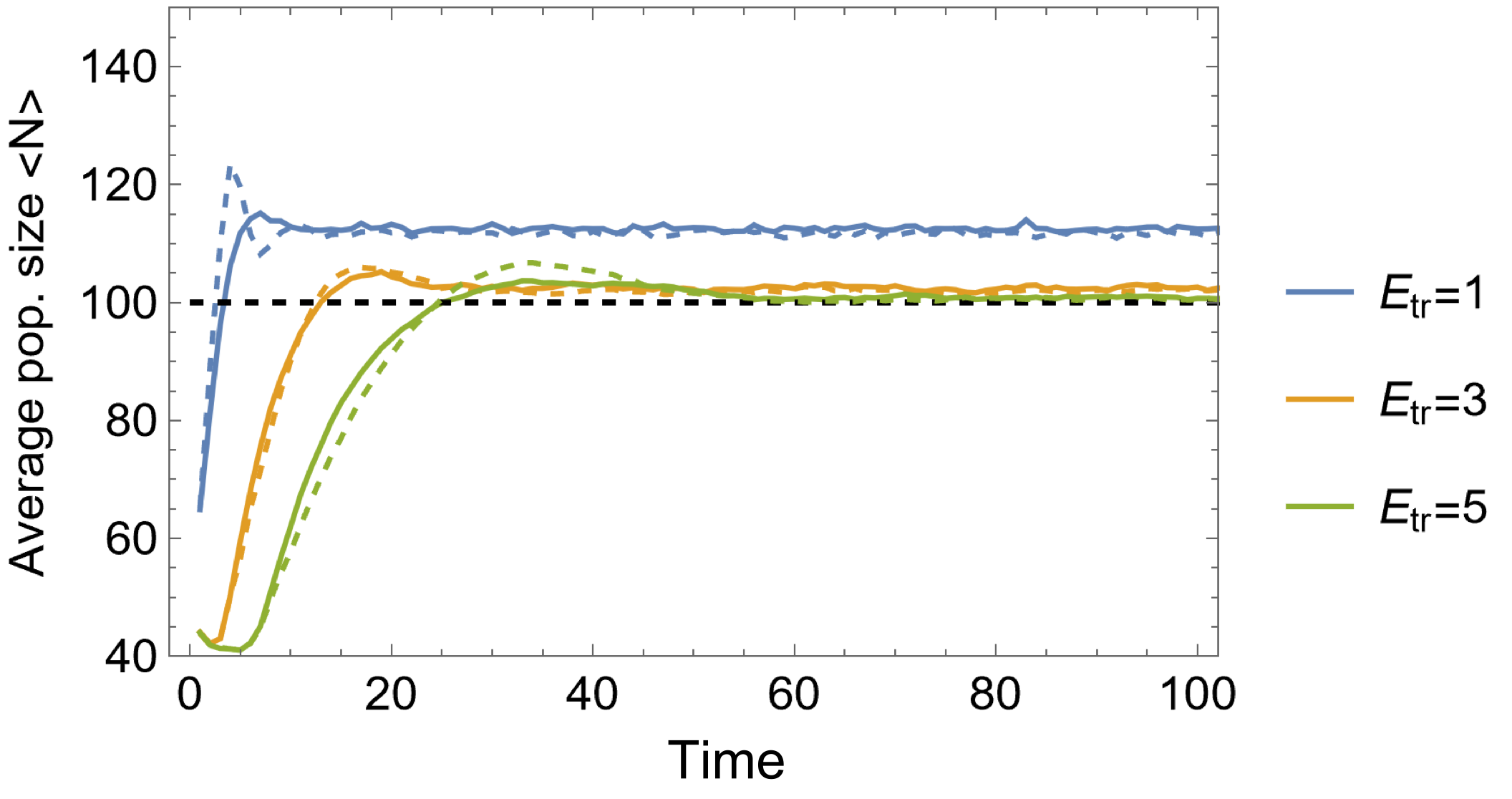

(c)	(d)

FIG.S1: **Growth of a protocell population depending on model parameters.**

The mean number of cells *< N*(*t*) *>* is obtained by averaging over 100 realizations with fixed initial conditions and model parameters.

(a) dependency of average number of randomly dividing protocells on housekeeping cost. Here, $p = 0.8,E_{tr} = 5, R = 30$.

(b) dependency of the average number of randomly dividing protocells on the probability of successful reproduction for $\Delta E = 0.3$. All other parameters are the same as in (a).

(c) dependency of the average population size of randomly or symmetrically dividing protocells on the reproduction threshold for $p = 0.7$and $\Delta E = 0.3$.

(d) dependency of the average population size of randomly or symmetrically dividing protocells on the reproduction threshold for $p = 1$(right), and $\Delta E = 0.3$. Solid curves denote random division, and dashed curves denote symmetrical division. The horizontal dashed lines show the estimates of the total population size for large *t* obtained via (A2).

The dependency of the average total number of protocells on the threshold is presented in Fig.S1c and Fig.S1d, for $p = 0.7$ and $p = 1$, respectively. Increasing the threshold value results in an increase of the average total population size for $p = 0.7$ and in a decreased population size for $p = 1$. Let us consider this phenomenon for symmetrically dividing cells. In the model, there are two causes of death of protocells: lack of resources and failed reproduction. In the case of high threshold values and small probability $p$, both daughter cells will be better prepared for the lack of resources in the next rounds (resources are proportional to $E_{tr}$ at cell birth). Hence they will mostly die due to failed reproduction. For small threshold values, the daughter protocells are vulnerable to the shortage of resources; indeed, in the considered case, the progeny will die in the absence of resources for two rounds in a row. In contrast, when the probability of successful reproduction is high ($p\approx1$), there is no loss of resources due to unsuccessful reproduction. In this case, although protocells with small thresholds remain vulnerable to the lack of resources, they gain advantage due to faster reproduction, and therefore, the number of new protocells per unit resource is greater than in the case of large threshold values.

Further increase of the threshold values does not measurably alter the time dependency of the average total number of the protocells. In Fig.10c,d, the time dependency of the average size of the population is shown for both random and symmetrical division of the protocells. Symmetrical division is advantageous for the population because the resource balance is more narrowly distributed around $\frac{E_{tr}}{2}$ than in the case of randomly dividing protocells, for large $t$. Therefore, symmetrically dividing protocells are better prepared for the lack of resources than randomly dividing cells.

Let us discuss the case of stochastic death events also. In this case we assume that at each round there is a nonzero probability $v$ that the given protocell will die due to the other reasons than the metabolism. Now in the coarse grained description (A1) the right hand side will be

$N{\Delta B}_{av}\left( t \right)= R- N\Delta E- \lambda\left( 1- p \right)N B_{av}-\nu B_{av} N ,$ (A3)

where the last term corresponds to the loss of resources due to the stochastic death events. Now the total number of protocells in the equilibrium will be

$N^{*}=\frac{R(1+p)}{2 \Delta E+\nu E_{tr}}$	(A4)

(A4) is true for small $\nu\ll1$, where we can assume that the average resource balance due to the stochastic deaths can be approximated linearly, for $\nu\sim1$ (A4) fails.

Here the threshold value of the reproduction of protocells enters explicitly in the expression of the total number of protocells in equilibrium, in contrast to (A1). Indeed, increasing the threshold value of reproduction of the protocells will decrease the equilibrium size of the population, since the probability that the given cell will die before reproduction is greater for the higher values of threshold, for the given probability of death events $v$.

***Appendix B. Competition between two populations of protocells***

The above introduced coarse grain model describing the average resource balance of the population is not applicable in case of the competition of two populations. Nevertheless, for a large difference between threshold values $E_{tr 2} >> E_{tr 1}$_,_ the simple model may still be used. Here we assume that two populations differ only by the threshold values, all other parameters are the same for both populations and there is no causes of death other than the metabolism $\nu=0$.

Indeed, for the large difference in the threshold values, it can be assumed that the total number of the population with large threshold value remains constant $N_{2} \approx const$. Now, we can use (1) for the first population, again assuming that the cells are identical. For large *t* and *E_tr_*_1_ *>* 1 the resource for balance of the first population is given by

$N_{1}{\Delta B}_{av 1}=N_{1}\frac{R}{N_{1}+N_{2}}-N_{1}\Delta E-\lambda_{1}(1-p)N_{1}, (B1)$

where ${\Delta E}_{1} = {\Delta E}_{2} = \Delta E$ and $p_{1}=p_{2}=p$. The first term in the right is the resource obtained by the first population. We assume that $\lambda_{1}=\frac{R/(N_{1}+N_{2})}{E_{tr}}$, that is, reproduction of the cells in a population depends on the sum of the sizes of the two populations and by the threshold value of the given population.

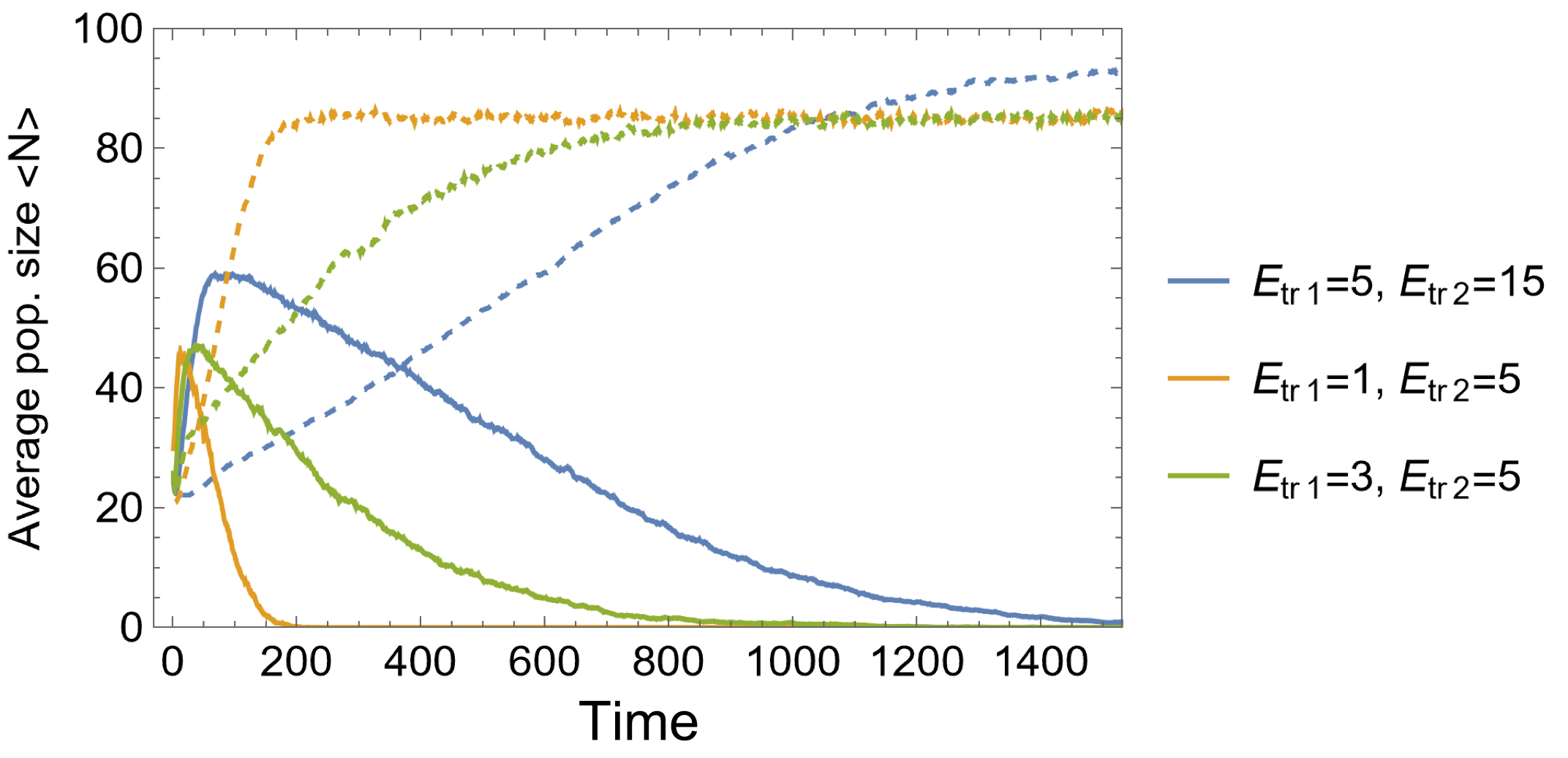

FIG.S2: **Competition between two protocell populations with different resource threshold values**. The solid and dashed lines represent average numbers of the cells with small and large threshold values, respectively. The other parameters are $\Delta E = 0.3, p = 0.7 R = 30$.

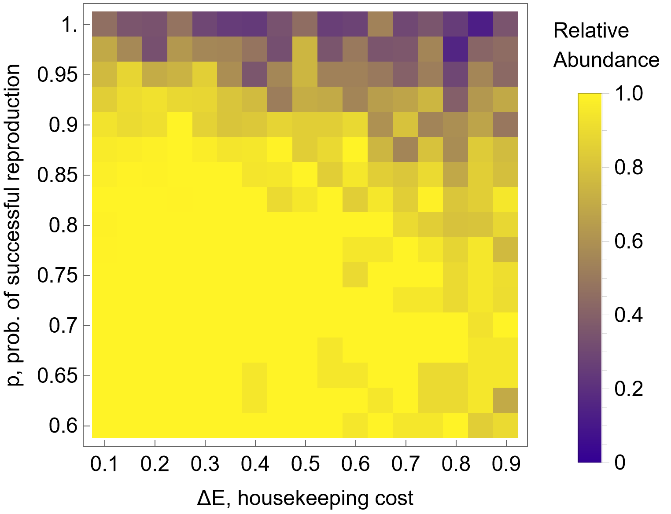

FIG.S3: **Competition of symmetrically dividing cells against randomly dividing cells for different values of successful reproduction probability** $\boldsymbol{p}$**and housekeeping cost** $\boldsymbol{\Delta}\boldsymbol{E}$. For the given value of $p\in\left[ 0.6, 1 \right]$ with step size $\delta p=0.025$, relative abundancy of symmetrically dividing cells is computed for $\Delta E\in[0.1,0.9]$ with step size equal to $\delta E= 0.05$. The color of each element represents the average (over $M=20$s independent simulations) fraction of symmetrically dividing cells in the whole (symmetric and random) population at time $T=5000$.

For the first population, we obtain $N_{1}^{*}+N_{2}=\frac{R(1+p)}{2\Delta E}$, that is, the final size of the first population is given by the same expression as in the absence of competition, except the initial presence of the second population $N_{2}\approx const$. Note that we assume that the second population is not reproducing until $N_{1}\to N_{1}^{*}$. Along the same lines, we obtain the resource increment for the second population, that is, $\Delta B_{2}\left( t \right)\propto\frac{2\Delta E}{1+p}-\Delta E-\lambda_{2}(1-p)B_{2}(t)$. From the last equation, it follows that if $p = 1$, the second population will not grow on average. Indeed, for $p = 1$, there is no loss of resources due to the failure in the reproduction process of cells in both populations. Therefore, the population that reproduces faster (a smaller threshold value) outcompetes its opponent. The competition outcomes for two populations with different threshold values are presented in Fig.S2. In all cases, the population with the smaller threshold value reaches the maximum first at approximately $\frac{R(1+p)}{2\Delta E}-N_{2}(0)$, but then starts declining in the course of the competition and loses to the population with the higher threshold value. As mentioned above, for $p<1$, symmetric division of resources between the daughter cells is advantageous over random division of the resources in the one population case (see Fig.S1c and Fig.S1d).

Symmetric division of the cells is similarly advantageous for two competing populations when $p_{1}=p_{2}=p<1$. Symmetrically dividing cells are better prepared to survive shortage of resources and therefore outcompete randomly dividing competitors with the same probability of successful reproduction $p_{1}=p_{2}=p<1$, see Fig.S3 where the average of the relative abundance of the symmetrically dividing cells in the environment $\frac{N_{sym}(T)}{N_{sym}(T)+N_{rand}(T)}$ is presented. The relative abundancies are calculated at $T=5000$. However, for $p_{1}=p_{2}=p\approx1$, the randomly dividing cells in average outcompete the symmetrically dividing counterparts. The point, is that for $p_{1}=p_{2}=p\approx1$ there is no loss of resources due to the failure in the reproduction events. The randomly dividing cells more likely produce asymmetrical progenies. The progeny with the larger resources will reproduce sooner, and hence will produce more progenies, than the progeny of a symmetrically dividing cell.

***Appendix C.***

***Deterministic description of (3).***

In the model, we introduced the elementary processes between the GE, in the absence of protocells

$*\underset{\to}{r}S, S+\mathcal{A+A}\underset{\to}{r_{\mathcal{A}}}3\mathcal{A,} S+\mathcal{A+N}\underset{\to}{r_{\mathcal{N}}}\mathcal{A}+2\mathcal{N}$ $\mathcal{A}\underset{\to}{d_{\mathcal{A}}}0, \mathcal{N}\underset{\to}{d_{\mathcal{N}}}0$

The propensities of each reaction is as follows $r, r_{\mathcal{A}} S\frac{m_{\mathcal{A}}\left( m_{\mathcal{A}}-1 \right)}{2}, r_{\mathcal{N}} Sm_{\mathcal{A}}m_{\mathcal{N}}, d_{\mathcal{A}}m_{\mathcal{A}}$and $d_{\mathcal{N}}m_{\mathcal{N}}$, where $m_{\mathcal{A}}$ and $m_{\mathcal{N}}$are the number of autonomous and non-autonomous GE, $S$ is the number of resource molecules available in the environment. To obtain the deterministic description of the process we assume that the number of all molecules is large $m_{\mathcal{A}},m_{\mathcal{N}}, S\gg1$. In this case, the time variation of the number of each type of molecules is given by the following dynamical system

$\frac{d S}{d \tau}=r-r_{\mathcal{A}}m_{\mathcal{A}}S-r_{\mathcal{N}}m_{\mathcal{A}}m_{\mathcal{N}}S$	(C1)

$\frac{d m_{\mathcal{A}}}{d \tau}= r_{\mathcal{A}}m_{\mathcal{A}}^{2}S-d_{\mathcal{A}} m_{\mathcal{A}}$	(C2)

$\frac{d m_{\mathcal{N}}}{d \tau}= r_{\mathcal{N}}m_{\mathcal{A}} m_{\mathcal{N}}S-d_{\mathcal{N}}m_{\mathcal{N}}$	(C3)

The dynamical system (C1-C3) have only one equilibrium state, that is autonomous-only GE in the environment given by $\left( S^{*},m_{\mathcal{A}}^{*},m_{\mathcal{N}}^{*} \right)=\left( \frac{{d_{\mathcal{A}}}^{2}}{r_{\mathcal{A}}r},\frac{r}{d_{\mathcal{A}}},0 \right)$, in general case. The coexistence of both types is possible only if $\frac{r_{\mathcal{N}}}{r_{\mathcal{A}}}=\frac{d_{\mathcal{N}}}{d_{\mathcal{A}}}$, a fine tuned condition which implies that there is effectively only one type of GE. In all other cases, the number of both types of GE tends to zero, while resources grow linearly.

Now, the condition of the stability of the autonomous-only equilibrium state can be found by analyzing the eigenvalues of the Jacobian matrix of (C1-C3) at that point, which has the following form

$J\left( S^{*},m_{\mathcal{A}}^{*},m_{\mathcal{N}}^{*} \right)=\left( \begin{matrix} -\frac{r_{\mathcal{A}} r^{2}}{{d_{\mathcal{A}}}^{2}} & -{2 d}_{\mathcal{A}} & -\frac{r_{\mathcal{N}}}{r_{\mathcal{A}}}d_{\mathcal{A}} \\ \frac{r_{\mathcal{A}} r^{2}}{{d_{\mathcal{A}}}^{2}} & d_{\mathcal{A}} & 0 \\ 0 & 0 & \frac{r_{\mathcal{N}}}{r_{\mathcal{A}}}d_{\mathcal{A}}-d_{\mathcal{N}} \end{matrix} \right)$

The eigenvalues are found from the following characteristic equation

$\left( \frac{r_{\mathcal{N}}}{r_{\mathcal{A}}}d_{\mathcal{A}}-d_{\mathcal{N}}-\lambda\right)\left( \left( -\frac{r_{\mathcal{A}} r^{2}}{{d_{\mathcal{A}}}^{2}}-\lambda\right)(d_{\mathcal{A}}-\lambda)+2\frac{r_{\mathcal{A}} r^{2}}{d_{\mathcal{A}}} \right)=0$	(C4)

From (C4) the eigenvalues are given by the following expressions

$$\left\{ \begin{aligned} \lambda_{1}=\frac{r_{\mathcal{N}}}{r_{\mathcal{A}}}d_{\mathcal{A}}-d_{\mathcal{N}} \\ \lambda_{2, 3}=\frac{-(a-d_{\mathcal{A}})\mp\sqrt{{(a-d_{\mathcal{A}})}^{2}-4{a d}_{\mathcal{A}}}}{2} \end{aligned} \right.$$

Where $a\equiv\frac{r_{\mathcal{A}} r^{2}}{{d_{\mathcal{A}}}^{2}}$. The autonomous-only equilibrium will be asymptotically stable if the reqal part of all eigenvalues will be negative $Re\left( \lambda_{1, 2,3} \right)<0$, which is only possible if

$$\left\{ \begin{aligned} \frac{r_{\mathcal{N}}}{r_{\mathcal{A}}}<\frac{d_{\mathcal{N}}}{d_{\mathcal{A}}} \\ r_{\mathcal{A}}r^{2}>{d_{\mathcal{A}}}^{3} \end{aligned} \right.$$

In the main text we focus on the case where $\frac{r_{\mathcal{N}}}{r_{\mathcal{A}}}>\frac{d_{\mathcal{N}}}{d_{\mathcal{A}}}$, which implies that the autonomous-only equilibrium is not stable, hence, a small variation from the equilibrium will result to the collapse of both types of GE.

***Appendix D.***

***Probabilities of the appearance of autonomous GE-only containing protocells due to reproduction***

In the main text, we provide the probabilities that the given daughter protocell will contain only autonomous GE (more than 1 element) (12) and (13) due to the reproduction of mother protocell, respectively for different reproduction scenarios (7) and (8).

We are interested in the given daughter protocell (let’s say the 1) will contain more than one autonomous element and no non-autonomous one, that is $m_{\mathcal{A}1}>1$ and $m_{\mathcal{N}1}=0$.

For the reproduction scenario (7) the calculation is straightforward.

Since, both types of genetic elements are pooled independently with uniform distribution, then we get

$P_{uni} \left( m_{\mathcal{A}1}>1,m_{\mathcal{N}1}=0 \right)=\frac{1}{m_{\mathcal{N}}+1}(1-\frac{2}{m_{\mathcal{A}}+1})$	(D1)

Where $m_{\mathcal{A}}$and $m_{\mathcal{N}}$ are the numbers of autonomous and non-autonomous genetic elements in the mother cell at the time of reproduction. The first multiplier in (D1) is the probability to choose 0 out of $[0,m_{\mathcal{N}}]$, since the probability of choosing any given number is $\frac{1}{m_{\mathcal{N}}+1}$. The second multiplier in (D1) is the probability to not choose 0 or 1 out of $[0,m_{\mathcal{A}}]$.

For the reproduction scenario (8), let us denote the amount of resources that the first daughter cell obtain due to the reproduction of the mother cell by $\pi\equiv\frac{B_{1}}{B}$. Now, $\pi$ is uniformly distributed in the interval $\left[ 0,1 \right].$ Now for any given $\pi$ the probability of $m_{\mathcal{A}1}>1,m_{\mathcal{N}1}=0$ is equal to

$P\left( m_{\mathcal{A}1}>1,m_{\mathcal{N}1}=0 | \pi\right)=\left( 1-\pi\right)^{m_{\mathcal{N}}}(1-\sum_{k=0}^{1} C_{m_{\mathcal{A}}}^{k}\pi^{k}{(1-\pi)}^{m_{\mathcal{A}}-k})$	(D2)

Where the first and second multipliers account for having no non-autonomous and more than one autonomous elements in the given daughter cell.

Now marginalizing over $\pi$ we will obtain the probability of $m_{\mathcal{A}1}>1,m_{\mathcal{N}1}=0$, that is

$P\left( m_{\mathcal{A}1}>1,m_{\mathcal{N}1}=0 \right)=\int_{0}^{1} P\left( m_{\mathcal{A}1}>1,m_{\mathcal{N}1}=0 | \pi\right) p\left( \pi\right)d\pi= = \frac{1}{m_{\mathcal{N}}+1}-\frac{1}{{m_{\mathcal{A}}+m}_{\mathcal{N}}+1}(1+\frac{m_{\mathcal{A}}}{{m_{\mathcal{A}}+m}_{\mathcal{N}}})$	(D3)

Subtracting (D3) from (D1) we find that

$\frac{1}{{m_{\mathcal{A}}+m}_{\mathcal{N}}+1}\left( 1+\frac{m_{\mathcal{A}}}{{m_{\mathcal{A}}+m}_{\mathcal{N}}} \right)-\frac{2}{{(m}_{\mathcal{A}}+1)(m_{\mathcal{N}}+1)}=$ $\frac{m_{\mathcal{N}}(m_{\mathcal{A}}({2m}_{\mathcal{A}}-1)+m_{\mathcal{N}}(m_{\mathcal{A}}-1)-1)}{({m_{\mathcal{A}}+m}_{\mathcal{N}}+1)(m_{\mathcal{N}}+1)(m_{\mathcal{A}}+1)(m_{\mathcal{A}}+m_{\mathcal{N}})}$	(D4)

(D4) is positive as long as the event $m_{\mathcal{A}1}>1,m_{\mathcal{N}1}=0$has non-trivial probability.

Under the symmetric reproduction scenario (9), the appearance of autonomous-only protocells due to the reproduction event is impossible if $m_{\mathcal{N}}>1$, that is at least two non-autonomous elements are present in the mother cell at the time of reproduction. Therefore, it is still possible due to the initial distribution of GE in the protocells and the death events as shown in the next section.

***Appendix E.***

***The impact of the initial distribution of GE in the appearance of autonomous-only protocells.***

The initial number of GE in protocells is given by the Poisson distribution with parameter$\mu$, same for all protocells for the given simulation,

$m_{0}^{i}=m_{\mathcal{A}0}^{i}+m_{\mathcal{N}0}^{i}=Poisson\left( \mu\right),i=1,\ldots N_{g0}$	(E1)

where $N_{g0}$ is the initial number of protocells containing GE (at the start of the first round of resource supply). The initial number of mutualists in each protocell is defined by a randomly (with uniform distribution) chosen integer from $\left[ 0,m^{i}\left( 0 \right) \right]$. Now, as greater $\mu$ as unlikely will be the appearance of autonomous only protocells, in general. Indeed, it is confirmed by observing the waiting times of the appearance of autonomous-only protocells.	Let us denote the time (measured by resource update rounds) at which the first mutualist-only protocell appears in the population by $\Theta$, the waiting time, which is a discrete random variable. If a mutualist-only protocell appears in the population due to the initialization (E1), then $\Theta=1$, that is, this protocell appears in the first round of resource supply. A mutualist-only protocell might not emerge at all during the simulation time $T$ (all GE can even die out during this time), in which case we assign $\Theta=T$. For convenience, we analyze the logarithm of the time of the first appearance of a mutualist-only protocell $Log\Theta$, with the ensemble average

$<Log\Theta> =\frac{1}{M}\sum_{l=1}^{M} Log\Theta_{l}$	(E2)

Here $M={10}^{3}$ is the total number of simulations, and $\Theta_{l}$ is the time at which the first mutualist-only cell appears in $l$th simulation. Under the worst-case outcome (for GE), when no mutualist-only protocells emerge in any of the *M* simulations, $<Log\Theta>=LogT$. Conversely, in the best-case outcome, a mutualist-only cell appears during the initialization of each simulation, $<Log\Theta>=0$.

a)
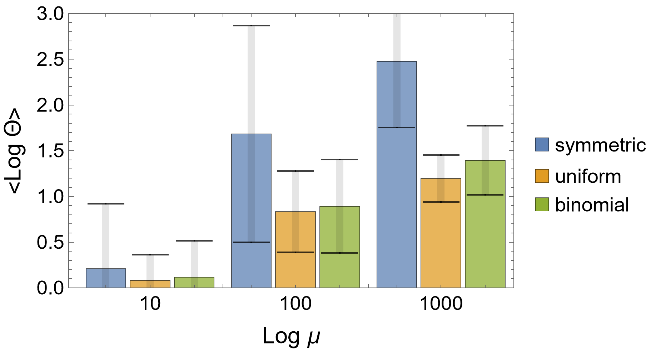
b)
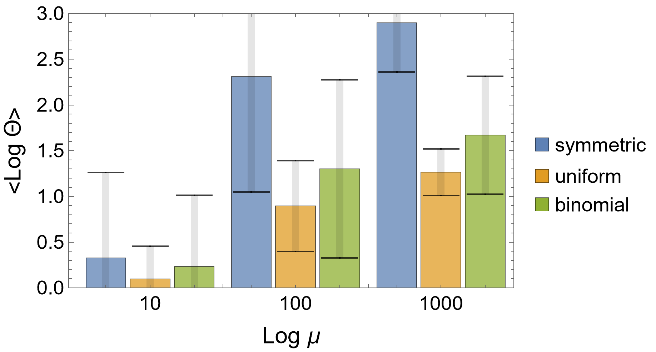

FIG. S4: **Kinetics of the appearance of mutualist-only protocells**. The labels of the charts are defined by the distribution of GE between daughter cells. The chart shows the dependency of the logarithm (base $10$) of waiting time of the appearance of mutualist-only protocells on the value of the parameter $\mu$ which describes the initial number of GE in the protocells (E1) for reproduction scenarios (7), (8) and (9). a) and b) subfigures show the simulation results for $r_{\mathcal{N}}\sim r_{\mathcal{A}}$ ($r_{\mathcal{N}}=0.0125, r_{\mathcal{A}}= 0.01$) and $r_{\mathcal{N}}\gg r_{\mathcal{A}}$ ($r_{\mathcal{N}}=1, r_{\mathcal{A}}= 0.01$), respectively. The simulation time is $T={10}^{3}$and the number of independent simulations is $M={10}^{3}$. The intervals show the standard deviation of the samples. The remaining parameters are as follows $d_{\mathcal{A}}=d_{\mathcal{N}}= 0.01, R = 30, \Delta E = 0.3, E_{c} = 0.01, p_{0} = 0.6, \mu= 100, K=1, E_{tr} = 5$.

The figure shows the dependency of the average waiting time (E2) and the standard deviation of the samples ($\sqrt{\frac{1}{M-1}\sum_{k}^{M} \left( Log\Theta_{l} -<Log\Theta> \right)^{2}}$) on the Poisson parameter $\mu$, the initial number of GE in each protocell (5), for the reproduction scenarios (7), (8) and (9), for both cases of $r_{\mathcal{N}}\sim r_{\mathcal{A}}$ and $r_{\mathcal{N}}\gg r_{\mathcal{A}}$. . Here $T={10}^{3}$, that, is the worst outcome is $<Log\Theta> = 3$. As expected, for the smaller initial number of GE ($\mu$) in each protocell, the likelihood of the appearance of a mutualist-only protocell is higher than for the larger $\mu$ in all scenarios, that is, the time of the first appearance of mutualist-only cells is shorter than it is for larger $\mu$. The ordering of the favorable reproduction scenarios is the same as for Fig.4, that is (7) is better than (8), which is better than (9), as it is expected from (D1) and (D3).
